# Strength of interactions in the Notch gene regulatory network determines patterning and fate in the notochord

**DOI:** 10.1101/2021.03.25.436857

**Authors:** Héctor Sánchez-Iranzo, Aliaksandr Halavatyi, Alba Diz-Muñoz

## Abstract

Development of multicellular organisms requires the generation of gene expression patterns that determines cell fate and organ shape. Groups of genetic interactions known as Gene Regulatory Networks (GRNs) play a key role in the generation of such patterns. However, how the topology and parameters of GRNs determine patterning *in vivo* remains unclear due to the complexity of most experimental systems. To address this, we use the zebrafish notochord, an organ where coin-shaped precursor cells are initially arranged in a simple unidimensional geometry. These cells then differentiate into vacuolated and sheath cells. Using newly developed transgenic tools together with in vivo imaging, we identify *jag1a* and *her6*/*her9* as the main components of a Notch GRN that generates a lateral inhibition pattern and determines cell fate. Making use of this experimental system and mathematical modeling we show that lateral inhibition patterning requires that ligand-receptor interactions are stronger within the same cell than in neighboring cells. Altogether, we establish the zebrafish notochord as an experimental system to study pattern generation, and identify and characterize how the properties of GRNs determine self-organization of gene patterning and cell fate.

## INTRODUCTION

Most of the information necessary to build an organism resides in its genome. The co-regulation of subsets of genes form gene regulatory networks (GRNs) that generate patterns of expression, which ultimately regulate cell fate and organ shape. Different types of GRNs regulate different patterning events. For example, some GRNs work in combination with gradients of morphogens to generate patterns at the embryo or organ scale (1). In contrast, other GRNs coordinate short-range interactions, generating self-organized patterns of gene expression at the cellular scale (2, 3). Understanding how different GRN topologies and the strength of their interactions regulate the generation of gene expression patterns constitutes a key challenge in developmental biology. However, research in this direction has been hindered by limited experimental systems that can be accurately modelled mathematically.

GRNs controlling short-range interactions produce diverse patterning events, such as lateral inhibition and lateral induction. Lateral inhibition involves a group of cells actively suppressing the expression of some genes in adjacent cells, thereby inducing them to adopt a different cell fate. In contrast, lateral induction involves cells inducing adjacent cells to adopt the same cell fate. Lateral inhibition and lateral induction patterns are two of the main patterns generated by Notch GRNs: one of the most representative signaling pathways that mediates local communication between cells. The Notch pathway is evolutionarily conserved and generates gene expression patterns that regulate cell fate decisions in a wide variety of organs (4–7). Signaling is triggered by interaction of a Notch receptor with a Notch ligand. Once they bind, the Notch intracellular domain (NICD) is cleaved and released inside the signal receiving cell. The NICD then translocates to the nucleus, where it activates Notch target genes (8).

The generation of either lateral inhibition or lateral induction patterns downstream of Notch has thus far been associated with different ligands. Lateral inhibition patterning has been described for the Delta-like (Dll) ligands and for Jag2 (9, 10) and generally occurs when Notch signaling activates the expression of a transcriptional repressor of the HES family that in turn inhibits the expression of the ligand in adjacent cells, preventing them from adopting the same cell fate (3, 11, 12). Mathematical simulations have shown that a lateral inhibition GRN can amplify small levels of noise in gene expression, leading to bi-stability and the generation of alternating patterns (13). Lateral induction has been shown for the ligand *Jag1*, whereby Notch activation by *Jag1* triggers the expression of the same ligand in the adjacent cells, promoting the same fate (14–16). It remains unknown whether lateral inhibition and lateral induction GRNs are restricted to specific ligands, or whether a given ligand can generate different patterns depending on the cellular and signaling context.

Other important parameters in a GRN are the nature and affinities of the ligand-receptor interactions. In the case of Notch, ligands can also interact with receptors in the same cell (17–19). This interaction, known as cis-inhibition, mutually inactivates both the ligand and receptor, and mathematical models have shown that it is required for patterning in the absence of cooperative interactions (20–22). Different ligands and receptors bind to each other in cis and trans with different affinities, and these affinities can be modulated by posttranslational modifications (3, 8). Altogether, these properties increase the complexity and diversity of Notch GRNs. For this reason, understanding how the topology and interaction parameters of these GRNs lead to pattern generation requires a combination of mathematical models and experimental systems that allow *in vivo* visualization and perturbation of Notch signaling components.

The notochord constitutes an underappreciated system that is ideal for studying the generation of Notch patterns. Initially, notochord coin-shaped precursor cells are arranged unidimensionally. These simple and well-defined cell-cell contacts greatly facilitate mathematical modeling and theoretical analysis, making it valuable for studying the relationship between GRNs parameters and patterns. In vertebrates, such as zebrafish, notochord precursors give rise to two different cell types (23): vacuolated cells, located in the inner part of the organ, that contain a large vacuole that provides hydrostatic pressure (24–26), and sheath cells, which form the surface of the cylindrical structure (23, 27) (Fig. 1A, Fig. S1 and movies S1, S2). The cell fate decision between vacuolated and sheath cells depends on Notch signaling (28). Inhibition of the Notch ligands *jag1a* and *jag1b* by morpholino injection leads to an excess in vacuolated cells, while overexpression of NICD promotes sheath cell fate (28). However, most of the components and topology of the GRN that coordinates cell fate in the notochord remain unknown.

**Figure 1.**
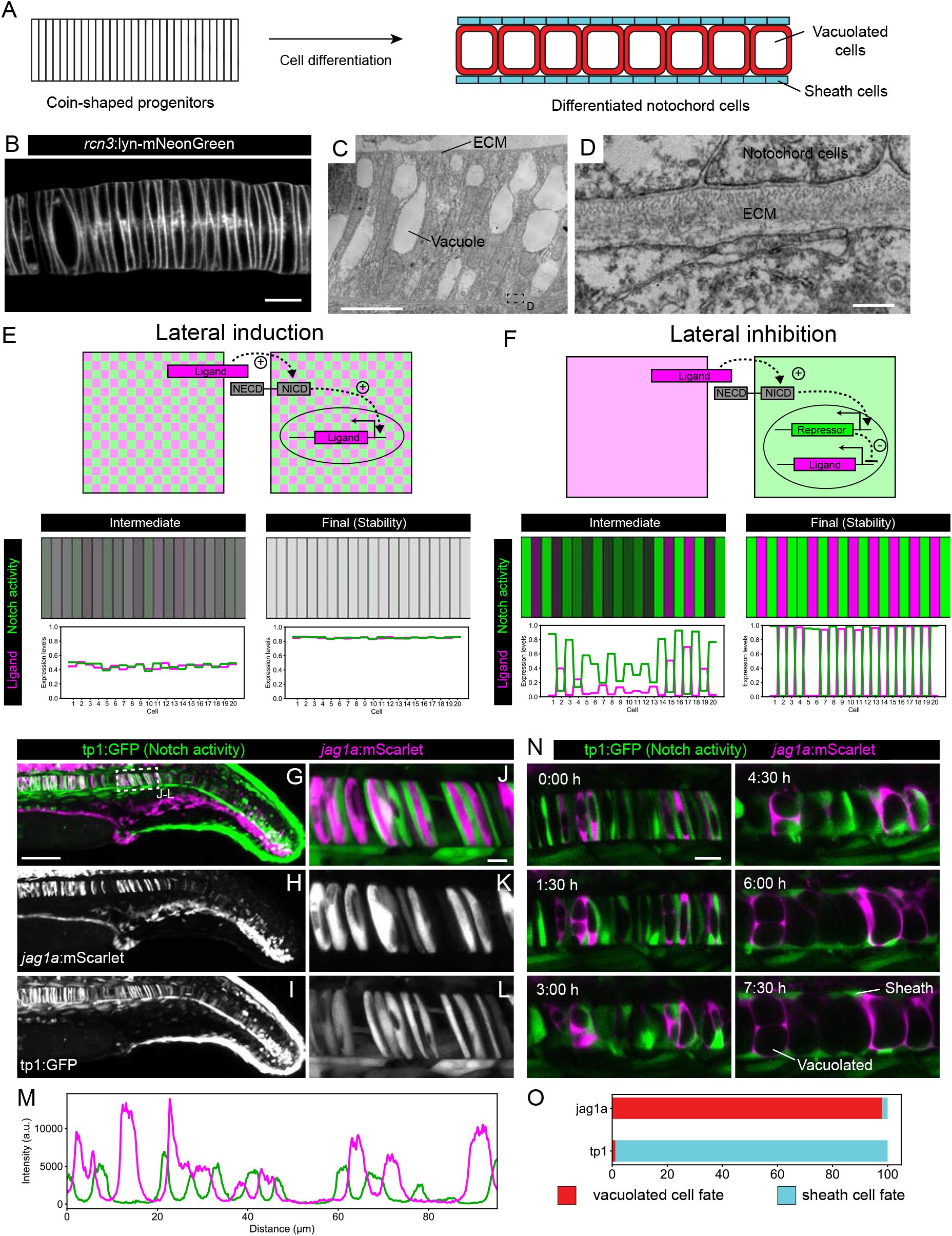
*Jag1a* generates a lateral inhibition pattern that correlates with fate. (**A**) Schematic representation of notochord development. At 18-19 hpf most of the notochord is composed by coin-shaped precursor cells. During the following 8 hours, progressively, in an antero-posterior order, coin-shaped precursors cells begin their differentiation into sheath cells and vacuolated cells. (**B**) Airyscan confocal section of a zebrafish notochord at 19 hpf using the *rcn3*:lyn-mNeonGreen transgenic line. (**C**) Transmission Electron Microscopy of a zebrafish notochord at 19 hpf. (**D**) Magnification of boxed area in (C). (**E**) (Top) Schematic representation of the model for a Lateral Induction Network shows a pair of cells where the ligand in one cell activates NICD release in the other cell. NICD activates ligand expression in its own cell. (Bottom) Representative simulation of this network applied to an array of cells unidimensionally arranged. (**F**) (Top) Schematic representation of the model for a Lateral Inhibition Network shows a pair of cells where the ligand in one cell activates NICD release in the other cell. NICD activates the expression of the repressor, which in turn inhibits Ligand expression. (Bottom) Simulation of this network applied to an array of cells unidimensionally arranged. (**G**-**L**) Maximal intensity projection of Airyscan confocal sections of a zebrafish tail at 22 hpf. (J-L) Magnification of boxed area in (G). n = 10 fish. **L**, Intensity profile across a horizontal line in panel (L). (**M**) *jag1a*:mScarlet and tp1:GFP expression levels across a 1 μm thick horizontal line on a single plane of the image shown in J. (**N**) Time lapse of optical sections of notochord cells using the tp1:GFP; *jag1a*:mScarlet double transgenic line. (**O**) Cell fate of cells expressing *jag1a* or the tp1:GFP at the coin-shape stage. Quantifications from images as shown in **N** (standard deviation jag1a = 2.696, tp1 = 2.631; n = 5 fish). Scale bars, 1 μm (D) 10 μm (B, C, J), 20 μm (N), 100 μm (G).

Here, we exploit the *in vivo* imaging and genetic manipulations that the zebrafish model offers to quantitatively study the generation of Notch patterns. Using the notochord experimental system, we show that *jag1a* generates a lateral inhibition pattern, a possibility thought to be restricted to the other Notch ligands (3, 29, 30). Using a combination of single-cell RNA-Seq analysis and genetic perturbations, we identify *her6*/*her9* and *jag1a* as the key genes that promote sheath and vacuolated fate. Our computational modeling further reveals that a stronger cis- than trans-inhibition is required for the generation of lateral inhibition patterns. We experimentally validate the role of cis-inhibition in our GRN, finding that *jag1a* is sufficient to disrupt the expression of Notch-target genes in the cells where it is expressed. Altogether, our results describe and characterize a novel Notch GRN that generates lateral inhibition patterns and determines cell fate.

## RESULTS

### 1. *Jag1a* and notch activity show a lateral inhibition pattern in the zebrafish notochord

Notch signaling generates patterns of gene expression by signaling at cell-cell contacts (31, 32). Thus, a prerequisite for the study of Notch patterning in the notochord is the characterization of cell-cell contacts. To describe the contacts between cells, we generated an *rcn3*:lyn-mNeonGreen transgenic line that labels the plasma membrane of all notochord cells. We observed that notochord precursor cells are coin-shaped and unidimensionally arranged one cell after another (Fig. 1B). Using transmission electron microscopy, we confirmed this cell arrangement and observed that coin-shaped notochord cells are isolated from the rest of the tissues by a layer of extracellular matrix (Figs. 1C, 1D). Thus, the contacts of each notochord cell are restricted to the two neighboring cells in the stack. This unidimensional geometry with very well-defined cell-cell contacts makes the notochord an ideal system to study Notch patterning.

Whether Notch signaling generates gene expression patterns in the notochord remains unknown. To understand the expression patterns that may be generated in this organ, we modeled lateral induction and lateral inhibition networks in the unidimensional arrangement of notochord cells. We first modeled a lateral inhibition network as a two component GRN, where the Notch ligand induces NICD cleavage in the adjacent cells, and NICD in turn induces ligand expression in the cells where it is located. This network gives rise to a homogeneous pattern, where all the cells have both high concentrations of NICD and ligand (Figs. 1E and S2A). Next, we modeled a lateral inhibition network (13). Here, the ligand also induces NICD cleavage in the adjacent cells, but in this case, NICD induces the expression of a repressor that in turn inhibits ligand expression. The result of this model is a NICD-ligand alternating pattern (Figs. 1F and S2B).

Then, we experimentally evaluated whether one of these two patterns was present in the notochord. The two zebrafish homologs of the mammalian *Jag1* – *jag1a* and *jag1b* – are the main Notch ligands in the notochord (Figs. S2C – S2F) (28). The non-homogenous expression of *jag1a* expression suggested that it might be involved in the generation of Notch patterns. To explore these patterns in high resolution, we generated a stable *jag1a*:mScarlet BAC transgenic line that recapitulates the endogenous *jag1a* mRNA expression (Figs. S2C-S2E), and crossed it to the tp1:GFP reporter of Notch activity (33). Unexpectedly, we found an alternating pattern (Figs. 1G – 1M) that resembles lateral inhibition, a pattern that has never been described for *Jag1*. Thus, our results show that *Jag1* is not restricted to the generation of lateral induction patterns as previously thought, but can also generate lateral inhibition patterns.

### 2. *Jag1a* and notch activity are early markers of notochord cell fate

Finding early markers of differentiation is important to understand cell fate decisions. However, no early marker of notochord cell differentiation has been reported to date. Having identified an alternating tp1-*jag1a* pattern, we evaluated whether it is associated with vacuolated and sheath cell fates. To test this, we used the tp1:GFP;*jag1a*:mScarlet double transgenic reporter, and followed notochord cells by time lapse *in vivo* imaging (Fig. 1N and movie S3). We found that *jag1a*-positive cells gave rise to vacuolated cells, while tp1-positive cells differentiated into sheath cells (Fig. 1O). Altogether, these results establish *jag1a* and tp1 as the first available markers of vacuolated and sheath cell fates.

### 3. *her9* and *her6* have a complementary expression pattern to *jag1a*

Having identified that the *jag1a*-Notch alternating pattern correlates with fate, we aimed to identify which are the components of the GRN that make possible this pattern. Notch lateral inhibition model predicts the presence of a Notch target gene that represses *jag1a* expression. This gene should have a mutually exclusive pattern with *jag1a*.

The bHLH genes of the HES/HEY families are good candidates as they are transcriptional repressors often activated by Notch signaling (34). In the notochord, *her9* has been shown to be a Notch downstream gene (28). However, the fact that no notochord phenotype was found for the *her9* knock down zebrafish (28), suggests a functional redundancy with other genes. To identify in an unbiased manner all the HES/HEY genes that repress *jag1a* we did single cell RNA-Seq analysis (35). We found that *her6* and *her9* are the most highly expressed genes of this family in the notochord at 18 and 24 hpf (Fig. 2A). To evaluate their expression pattern, we analyzed mRNA expression by fluorescent *in situ* hybridization based on a hybridization chain reaction (HCR). *her6* and *her9* were expressed in an alternating pattern with *jag1a* (Figs. 2B – 2M). In contrast, *her12*, which was expressed at a much lower level according to the RNA-Seq, was not detected in the notochord by HCR (Fig. S3). The observed alternating patterns suggest that *her6* and *her9* could repress *jag1a* expression in the notochord.

**Figure 2.**
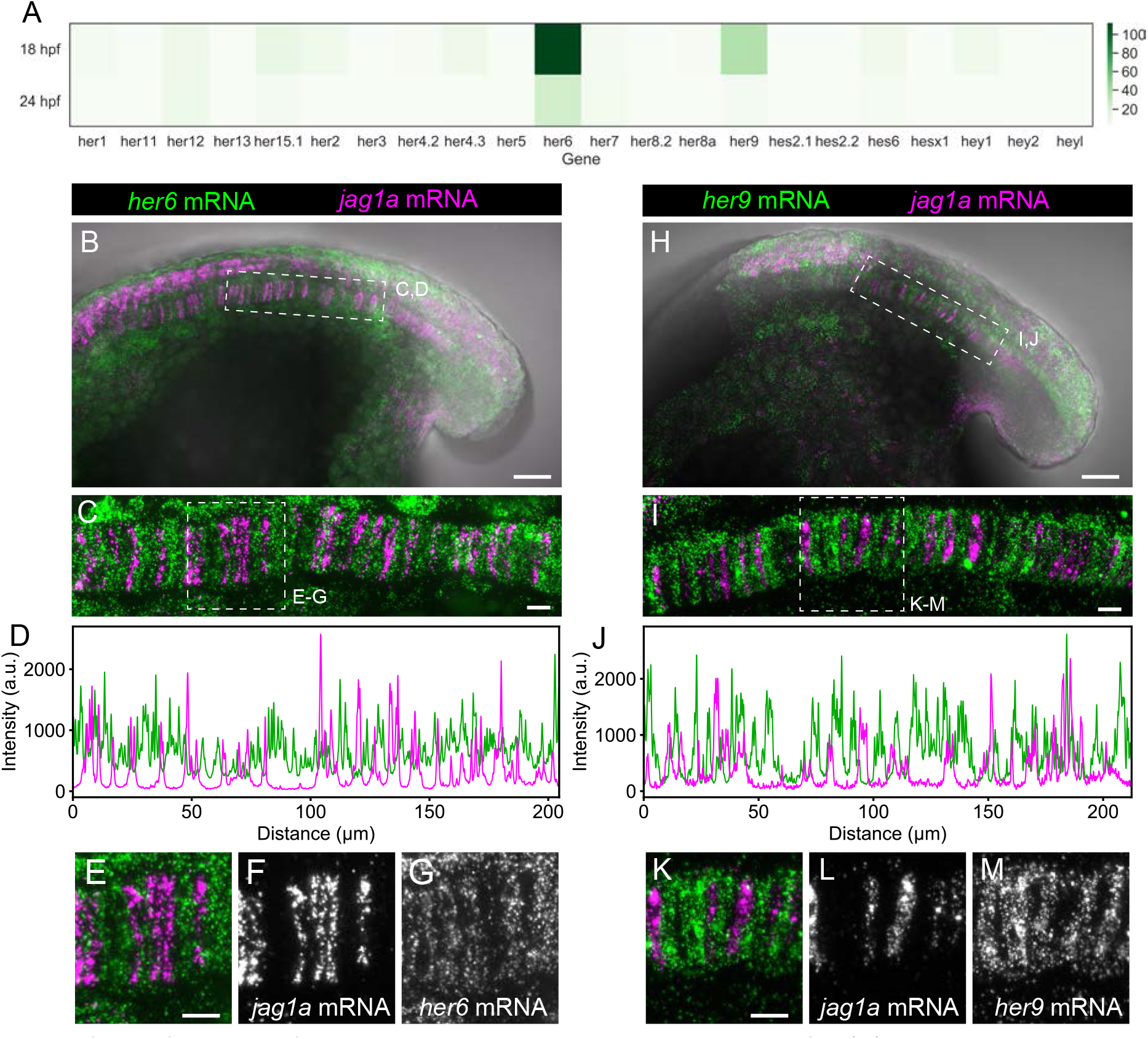
*her9* and *her6* show a complementary pattern to *jag1a*. (**A**) Heatmap showing the expression levels of the zebrafish HES/HEY family genes. Values represent average normalized UMIs in all notochord cells at 18 and 24 hpf. (**B**) Projection of confocal optical sections of 18 hpf zebrafish stained with in situ HCR probes against *her6* (green) and *jag1a* (magenta). Transmitted light is shown in gray scale. (**C**) Maximal projection of confocal Airyscan optical sections of the boxed area in (B)**. D**, Intensity profile of *her6* (green) and *jag1a* (magenta) along the notochord based on in situ HCR shown in (C). (**E**-**G**) Magnified views of boxed area in (C), n = 8. (**H**-**M**) Analogous images to (B-G) based on the *her9* probe instead of *her6* probe, n = 9. Scale bars, 50 μm (B, H), 20 μm (C, E, I, K).

Aside from the ligand and repressor, the other main component of a lateral inhibition Notch GRN is the Notch receptor. By single cell RNA-Seq data analysis (35) we found that *notch2* was detected in most cells at the highest levels at 18 and 24 hpf (Figs. S4A – S4F). *notch2* notochord expression was confirmed by fluorescent HCR (Figs. S4G – S4H). Altogether, we identified the main components of the lateral inhibition GRN, finding *her6* and *her9* as candidate genes to repress *jag1a* expression, and *notch2* as the main Notch receptor in the notochord.

### 4. *her6* and *her9* inhibit *jag1a* expression

To directly assess if *her6* and *her9* are sufficient to inhibit *jag1a* expression, we established notochord-specific genetic mosaics. To that end, we identified a highly specific notochord promoter to overexpress *her6* or *her9*, while simultaneously labeling the perturbed cells. Making use of the single-cell RNA-Seq dataset (35), we identified *emilin3a* as the gene that offers the best balance between notochord specificity and high expression levels. We cloned a 5 kb promoter upstream of the coding region and showed that it is sufficient to drive gene expression in the notochord in a highly specific manner (Fig. S5). Next, we used the identified promoter and the p2a system (36) to generate *her6* or *her9* gain-of-function cells concomitantly with GFP expression, or only-GFP as a control. For each of these constructs, we quantified the level of *jag1a*:mScarlet expression in the GFP-p2a-*her6*, GFP-p2a-*her9* or only-GFP positive cells in comparison to the rest of the notochord. We found that GFP-p2a-*her6* and GFP-p2a-*her9* cells had a lower level of *jag1a*:mScarlet than only-GFP cells, indicating that *her6* and *her9* repress *jag1a* expression in a cell autonomous manner (Fig. 3A-G). This result was confirmed by quantifying endogenous *jag1a* mRNA expression by fluorescent HCR (Fig. S6).

**Figure 3.**
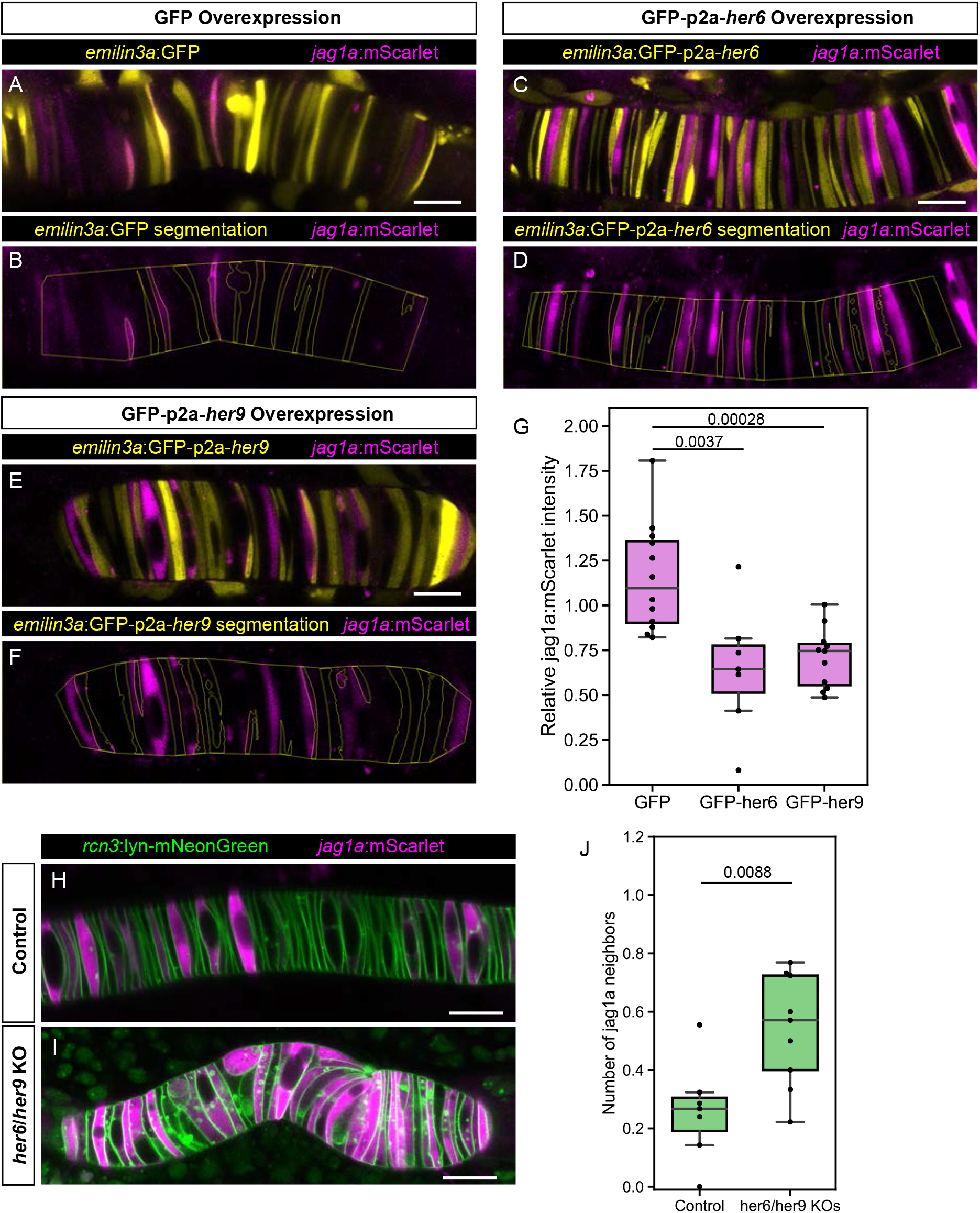
*her6* and *her9* inhibit *jag1a* expression. (**A** - **E**) Airyscan confocal optical sections of live 22 hpf transgenic *jag1a*:mScarlet zebrafish injected with *emilin3a*:GFP (A and B), *emilin3a*:GFP-p2a-*her6* (C and D) or *emilin3a*:GFP-p2a-*her9* (E and F). DNA constructs were injected at the 1-cell stage together with I-SceI protein. (B, D, F) show the boundary of GFP segmentation in A, C and E, respectively, and manual outline of the notochord. **G**, Quantification of *jag1a*:mScarlet intensity inside GFP-positive cells segmented as exemplified in (B, D, F). Values in the plot represent the intensity of *jag1a*:mScarlet inside segmented cells divided by the *jag1a*:mScarlet intensity inside the notochord outside of the segmented cells. Each point represents an individual fish. Two-tailed p-values are shown in the plot. **<u>H</u>**, Airyscan confocal sections of embryo at 22 hpf injected with only Cas9 (H) or Cas9 and her6/her9 gRNAs (I). **J** Quantification of the average number of jag1a-positive cells directly adjacent to each jag1a-positive cell. Each individual point in the plot represents the average value for an independent fish. Two-tailed p-value is shown in the plot. Scale bars, 20 μm.

Having identified *her6* and *her9* as genes sufficient to inhibit *jag1a* expression, we studied if these genes are necessary for lateral inhibition patterning in the notochord. To this end, we generated her6/her9 double transient knock-outs in a *jag1a*:mScarlet;*rcn3*:lyn-mNeonGreen background, and quantified the number of *jag1a*-positive cells that are found adjacent to each *jag1a*-positive cell. We found this value to be increased upon *her6* and *her9* gene deletion, showing that *her6* and *her9* are necessary for lateral inhibition (Fig. 3H-J). Altogether, we show that *her6* and *her9* are the repressors in the GRN that generate a lateral inhibition pattern in the notochord.

### 5. *her6*/*her9* and *jag1a* determine notochord cell fate

To test if the identified GRN genes are sufficient to determine cell fate, we first expressed GFP-p2a-*her6*, GFP-p2a-*her9* or only-GFP in a mosaic fashion in the notochord cells, and evaluated its effect on cell fate. At 2 days post-fertilization (dpf), a stage where vacuolated and sheath cells can be distinguished, we found a lower proportion of vacuolated cells in GFP-p2a-*her6* and GFP-p2a-*her9* expressing cells. This result indicates that *her6* and *her9* are sufficient to determine sheath cell fate (Fig. 4A – 4D).

**Figure 4.**
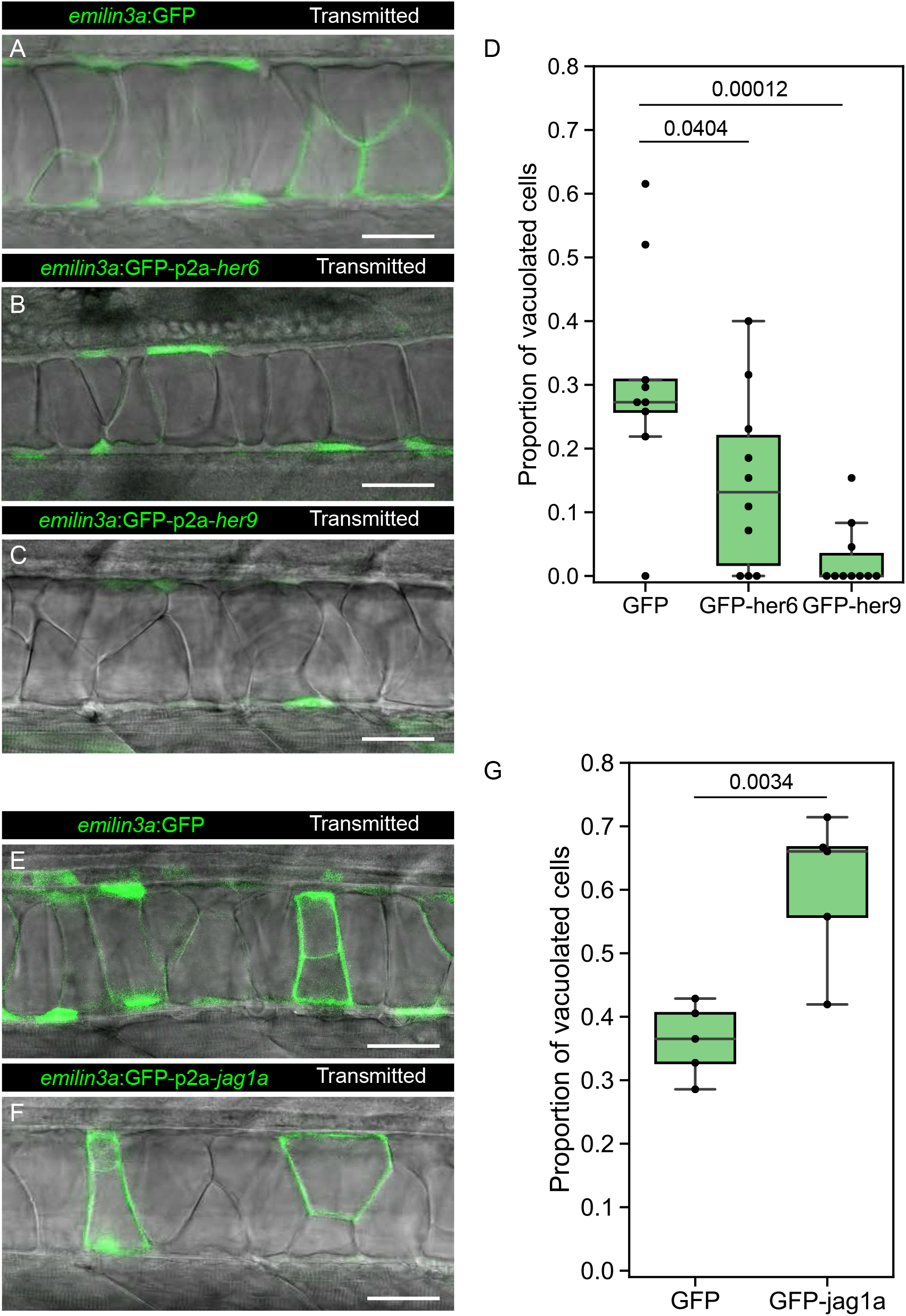
*her6*, *her9* and *jag1a* determine cell fate in the zebrafish notochord. (**A**-**C, E-F**) Confocal optical sections of 2 dpf live zebrafish that were injected with the *emilin3a*:GFP (A, E), *emilin3a*:GFP-p2a-*her6* (B), *emilin3a*:GFP-p2a-*her9* (C) or *emilin3a*:GFP-p2a-*jag1a* (F) constructs. DNA constructs were injected at the 1-cell stage together with I-SceI protein. (**D** and **G**) Proportion of vacuolated cells at 2 dpf are shown. Each point in D, G represents an independent fish quantified from on z-stack confocal planes. 2-tailed p-values are shown in D and G. Scale bars, 50 μm.

Next, we expressed GFP-p2a-*jag1a* or only-GFP. Interestingly, we found that the Notch ligand *jag1a* is sufficient to drive vacuolated cell fate in the same cells where it is expressed (Fig. 4E – 4G). Taken together, our results show that not only the Notch targets *her6*/*her9* drive cell fate, but also the Notch ligand *jag1a* determines cell fate on the same cell where it is expressed.

### 6. Stronger cis than trans interaction are required for lateral inhibition patterning

After observing that *jag1a*, a Notch ligand, drives vacuolated cell fate on the same cell where it is expressed, we next investigated the mechanism mediating this process. First, we explored a potential signaling role of the ligand intracellular domain. It has been shown that upon Notch-ligand trans-interaction, not only the NICD is cleaved in the receiver cell, but also the intracellular domain of some ligands, including Jag1, is cleaved inside the sender cell, leading to bidirectional signaling (37–42). The intracellular domain of *jag1a* (JICD) would then inhibit Notch signaling in the sender cell (38). Thus, overexpression of the full-length ligand in our experiment would increase the amount of ligand that is available to be cleaved, leading to Notch inhibition and promoting vacuolated cell fate. To test this hypothesis, we expressed mScarlet-p2a-JICD or only-mScarlet in a mosaic fashion in notochord cells. We did not observe any effect of JICD on cell fate (Fig. S7), showing that JICD signaling is not sufficient to explain the *jag1a* effect on fate in the notochord.

Next, we considered two different signaling circuits that could explain how *jag1a* can promote vacuolated cell fate in the cells where it is expressed. First, through trans-interactions with the Notch receptor, *jag1a* could activate Notch signaling and as a consequence, *her6*/*her9* expression in their neighbors. *Her6* and *her9* would inhibit *jag1a* in the neighbors, and this would in turn diminish the amount of Notch signaling that the initial cell receives and promote vacuolated cell fate. A second possible explanation comes from the observation that when Notch ligands are expressed in the same cell as the Notch receptor, they can mutually inhibit each other through cis-inhibition. Thus, overexpression of *jag1a* would deplete the Notch receptor in a cell-autonomous manner, making this cell non-responsive to Notch signaling, and thus, promoting vacuolated cell fate (Fig. 5A).

**Figure 5.**
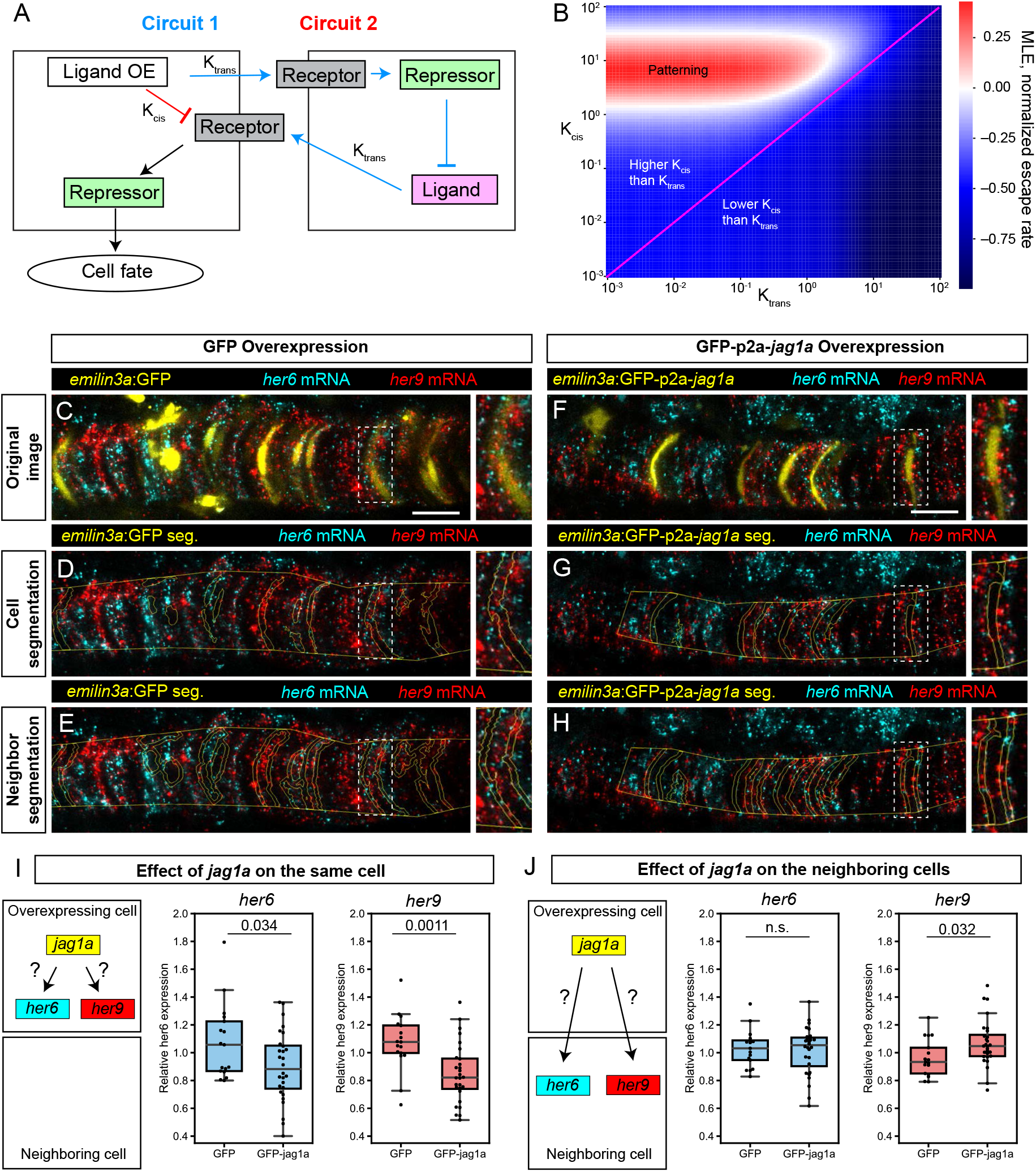
Modeling and experimental results of *cis* and *trans* interactions in the notochord. (**A**) Two possible circuits may explain the effect of *jag1a* on fate on the cell where *jag1a* is overexpressed. Circuit 1 is based on a possible role of cis-inhibition of the notch receptor by the ligand. Circuit 2 is based on the interaction of ligand and receptor in trans. Cells where we overexpress the ligand is represented as the cell on the left. Adjacent cells are represented on the right. OE, overexpression. (**B**) Escape rates from the homogeneous steady state (indicated by Maximum Lyapunov Exponents, or MLE) as a function of Kcis and Ktrans parameters. Positive MLE values (red) support patterning, while negative MLE values (blue) do not. (**C** - **H**), Airyscan confocal planes of fixed 22 hpf transgenic injected with *emilin3a*:GFP (C-E) or *emilin3a*:GFP-p2a-*jag1a* (F-H) constructs. GFP was detected by antibody staining and *her6* and *her9* mRNA by *in situ* HCR in whole mount embryos. (D and G) show the notochord outline manually selected and the outline of GFP-positive cells automatically segmented. (E and H) show the outline of the manually selected notochord and the neighborhood to the GFP-positive cells. On the right side of each panel, a magnified view of the boxed region is shown. (**I**, **J)** Quantification *her6* and *her9* mRNA expression after GFP-based segmentation as shown in (D, G) or (E, H), respectively. Values of *her6* and *her9* expression levels inside the segmented area inside the notochord were divided by the expression levels of the same genes in the region outside the segmented area, also inside the notochord. Each point represents a different fish. Two-tailed p-values are shown in the plots. n.s., non-significant. Scale bars, 20 μm.

To test the relative contribution of each of these circuits in patterning the notochord and in cell fate, we implemented a mathematical model that includes ligand-receptor interactions both in trans – between neighboring cells – and in cis – within the same cell – based on Sprinzak *et al* (21) (Fig. S8). Receptor-ligand cis and trans interactions are represented by the K_cis_ and K_trans_ parameters, respectively. Next, we used this model to dissect which combinations of cis and trans interactions lead to the observed lateral inhibition pattern (Fig. 1 G-M). To do so we evaluated the stability of the homogeneous steady state (HSS) depending on K_cis_ and K_trans_. The HSS is defined as the steady state where all the cells have identical concentrations of Notch ligand, receptor and repressor. When the HSS is stable, the system remains in this homogenous state and no patterning occurs. HSS stability can be evaluated by performing linear stability analysis to calculate the Maximal Lyapunov Exponent (MLE), which represents the exit speed from the homogeneous steady state. Thus, a positive MLE represents an unstable HSS, and this leads to patterning. We found that in the absence of cooperativity, patterning only occurs in a region of the parameter space where K_cis_ is higher than K_trans_ (Fig. 5B). This suggests that the observed lateral inhibition pattern not only requires the presence of Notch-ligand cis-interactions, but also, that such cis-interactions are stronger than trans interactions.

Our modeling predicted that the cis-circuit should be more active than the trans-circuit in the notochord. To test this prediction experimentally, we expressed GFP-p2a-*jag1a* or only-GFP in a mosaic manner in the notochord, and quantified the effect on *her6* and *her9* expression both within the same cell and in the neighboring cells. Interestingly, we found only a minor or no increase in *her6*/*her9* expression in the neighboring cells (Figs. 5C, 5D, 5F, 5G, 5I), suggesting a small Notch-ligand trans-interaction. However, we observed a strong reduction of *her6*/*her9* expression in the *jag1a*-expressing cells (Figs. 5E, 5H, 5J), indicating a strong cis-effect of *jag1a* expression. These results show that *jag1a* expression has a stronger effect on its own cell than in its neighbors, and supports the prediction of our model, where we showed that a stronger cis than trans interaction is required for the generation of lateral inhibition patterns.

## DISCUSSION

The unidimensional arrangement of cells in the zebrafish notochord, combined with its binary cell fate decisions, make it a unique model to study the properties of the Notch GRN that determines its patterning. One of the most important genetic interactions in a Notch GRN is how the expression of the ligands is regulated by Notch signaling. Previously, it was generally accepted that Notch signaling activates *Jag1* expression leading to lateral induction patterns (3, 29, 30). Here we show that Notch signaling, through the activation of the transcriptional repressors *her6* and *her9*, inhibits *jag1a* expression in the notochord, leading to the generation of lateral inhibition patterns. Importantly, *Jag1* is expressed in many other tissues apart from the notochord, including heart, inner ear, muscle and kidney (43–46), suggesting that the identified GRN may be relevant for pattern generation in these other contexts.

Another key part of a Notch GRN that may affect patterning, is whether upon ligand-receptor interaction, there is unidirectional or bidirectional signaling. In the bi-directional signaling situation, not only the cell expressing the receptor would receive a signal, but also the cell expressing the ligand. This signal would be mediated by the intracellular domain (ICD) of the ligand. However, the role of ligands ICDs remains unclear. Previous work showed that the ICD of JAG1 and DLL1 modulate cell differentiation, proliferation and Notch signaling (37–41). In contrast, other studies found little or no effect of DLL1-ICD, DLL4-ICD and JAG1-ICD on gene expression and migration in endothelial cells (42). In agreement with the latter, we found no role of the zebrafish *jag1a*-ICD on cell fate. Further research will be needed to elucidate if the role of ligands ICDs depends on the signaling context, and whether different cell types respond differently to ICDs.

Patterning not only depends on the topology of a GRN, but also on the strength of each of the interactions. Here, using mathematical simulations supported by experimental results, we shed light on which combinations of parameters promote pattern generation. Specifically, we find that a stronger Notch-ligand interaction in *cis* than in *trans* is key for pattern generation. Importantly, this does not mean that trans-interactions are not needed. In absence of such interactions, there would be no communications between cells and thus no lateral inhibition patterning.

The strength and signaling efficiency of *cis* and *trans* interactions in Notch GRNs depend on the specific ligand-receptor pair (3, 47–49). Some DLL, such as DLL4, activate Notch signaling in trans more strongly than Jagged ligands (47). On the other hand, the *Drosophila* homolog of Jagged genes, *serrate*, inhibits Notch receptors in cis more efficiently than Delta ligands (19, 50–52). The possibilities of imaging and genetic manipulation that the zebrafish offers, together with the unique cell-cell contacts in the notochord, will make this organ a very valuable *in vivo* system to evaluate the properties of not only endogenous ligands, but also other Notch ligands, to better understand how cis and trans parameters determine pattern generation.

Our results not only explain how Notch drives pattern generation, but also how cell fate is determined during notochord development. We identified Notch activity, and its downstream genes *her6* and *her9*, as key determinants of sheath cell fate in the notochord. In some tissues, including skeletal muscle, intestine and neural systems, a higher Notch activity is related to stemness, while a lower Notch activity is related to differentiation (53–57). This raises the interesting hypothesis of whether sheath cells can be considered as only partially differentiated notochord cells. In agreement with this concept is the recent finding that upon vacuolated cell damage, sheath cells develop vacuoles and partially restore notochord structure (58, 59). However, a possible role of Notch signaling during notochord regeneration is yet to be tested. Several pieces of evidence suggest that the GRN that we have identified is not exclusive to zebrafish. Previous studies based on BAC transgenesis showed that *Hes1*, the mammalian homolog of *her6* and *her9*, is expressed in the mouse notochord, suggesting it may play a role in the patterning of the mammalian notochord (60). Problems in notochord development have been associated with defects in spine morphogenesis (61–64). Interestingly, mutations in JAG1 and NOTCH2 (65, 66), the human homologs of the main ligands and receptor in the zebrafish notochord, lead to vertebrae malformations in human Alagille Syndrome. This suggests that spine problems in this human syndrome may be the result of defective Notch patterning during notochord development. Thus, in this study we describe a GRN that is likely conserved across vertebrates, opening the door to better understand how mutations in *JAG1* or *NOTCH2* lead to the problems observed in the human disease.

In non-vertebrate chordates such as ascidians, a single cell type performs the two main functions of both sheath cells and vacuolated cells: covering the surface and producing the fluid (67, 68). From an evolutionary perspective, it is plausible that Notch signaling was involved in dividing these possible ancestral functions into two different cell types. We speculate that Notch- or Hes-responsive enhancers were co-opted during vertebrate evolution to control the expression of the key genes necessary for vacuolated and sheath cell functions, making possible the specialization of the two different cell types. Given how frequently Notch signaling determines cell fate across development, Notch could represent a general mechanism that facilitated division of functions between different cells, promoting the evolution of new cell types.

Altogether, we have established the notochord as a new model system to study the principles that determine the pattern generation. Using a combination of mathematical modeling, single cell RNA-Seq analysis and genetic perturbation approaches, we identified *jag1a*, *her6*, *her9* and *notch2* as the key genes that determine cell fate and patterning. We expect that the GRN properties identified in this study will help understand the principles underlying patterning and cell fate decisions across multicellular organisms.

## Supporting information

Plasmid sequences

## METHODS

### Animal handling and generation of transgenic lines

The construct to generate Tg(*jag1a*:mScarlet) transgenic line was generated by BAC recombineering using the BAC CH211-21D8. We first used EL250 (*70*) bacteria to recombine first the iTol2Amp cassette ( (*71*), primers 1 and 2, Table S1) and substitute the loxP site in the BAC backbone. To recombine the mScarlet sequence into the BAC, we first used Gibson Assembly to substitute mCherry-p2a-CreERT2 by mScarlet in the mCherry-p2a-CreERT2-loxP-kan-loxP plasmid (*72*) to generate an mScarlet-FRT-kan-FRT plasmid (Data S1). Then, we used the primers 3 and 4 (Table S1) to amplify and recombine the mScarlet-FRT-kan-FRT into the ATG of jag1a in the BAC CH211-21D8. Finally, we removed the kanamycin resistance by activating flipase expression in the EL250 bacteria.

To clone the *emilin3a*:mScarlet plasmid (Data S2) we selected the 5 kb upstream of the *emilin3a* promoter and cloned it upstream of mScarlet in a tol2 plasmid. The *rcn3*:lyn-mNeonGreen construct (Data S3) was generated by Gibson Assembly using the previously described *rcn3* promoter (*24*).

*jag1a*:mScarlet, *emilin3a*:mScarlet and *rcn3*:lyn-mNeonGreen were injected at the one cell stage using tol2 transposase. To establish the stable transgenic lines, we crossed the fish by wild type until we found 50% of the progeny transgenic, indicative of a probable single insertion. For the *rcn3*:mNeonGreen transgenic line, due to the high variability in gene expression between different lines, we selected the most notochord specific line among 5-10 different founders.

**Table S1.**
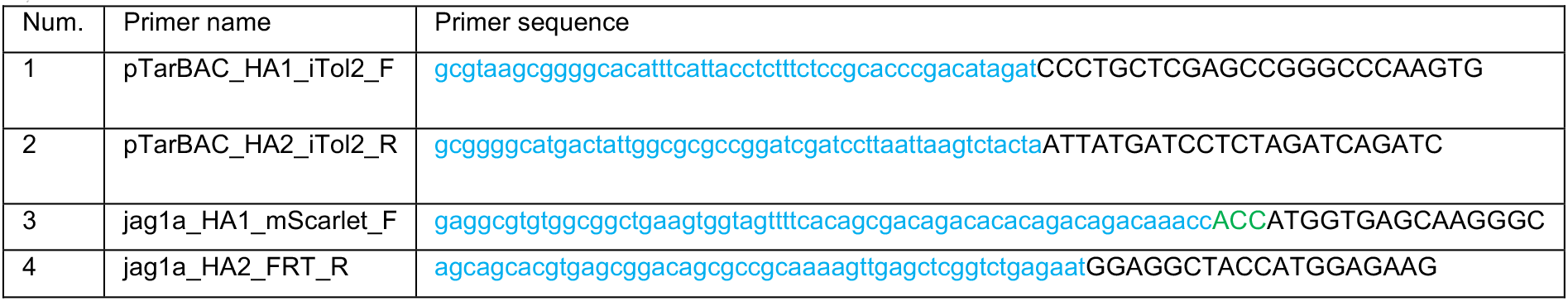
Primers 1 to 5 were used for cloning of the DNA constructs used to generate the transgenic zebrafish lines. Red, homology arms. Green, minimal Kozak sequence. Capital letters, overlapping with template sequence. F, forward; R, reverse.

### *her6* and *her9* Knock-out

To generate *her6* and *her9* transient knockout (crispants), we designed guide RNAs (gRNAs) targeting the beginning and the end of both *her6* and *her9,* resulting in whole gene deletion. Guides were identified using CRISPRscan (*73, 74*) and synthesized as previously described (*75*) (Primers 5-9, Table S2). Primers 10-13 (Table S2) were used for the detection of the deleted allele. This allele was found in all the embryos displaying a shortened axis (10/10). Only embryos showing this phenotype were used for further analysis by imaging. The injection mix included custom-produced Cas9-GFP at 2.4 mg/mL, KCl 300 mM and the four gRNAs, each of them at 12.5 ng/μL. Embryos where Cas9 but no gRNA was injected, were used as a control. Heterozygous embryos for both *rcn3*:mNeonGreen and *jag1a*:mScarlet transgenes were used in this experiment. Cells with *jag1a*:mScarlet intensity lower than 10% of the maximum intensity value in each image were considered negative for *jag1a*.

**Table S2.**
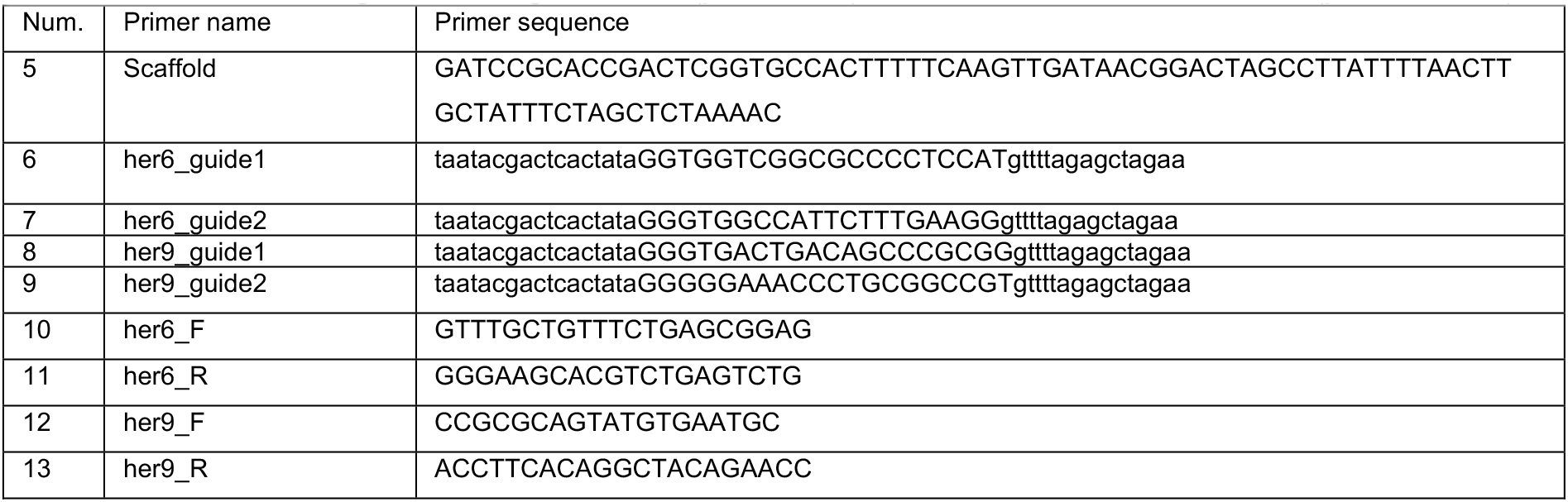
Primers used to generate the guide RNAs (primers 5-9) and detection of the deleted allele (primers 10-13).

### Cell fate analysis

*emilin3a*:GFP (Data S4), *emilin3a*:mScarlet (Data S2), *emilin3a*:GFP-p2a-*her6* (Data S5), *emilin3a*:GFP-p2a-*her9* (Data S6) or *emilin3a*:mScarlet-p2a-*jag1a* (Data S7) were cloned using Gibson Assembly using as template synthesized *her6*, *her9* and *jag1a* cDNAs. These plasmids were injected at the one cell stage using Isce-I as previously described (*76*). GFP fluorescence and transmitted light were imaged in vivo at 2 dpf. Quantifications were made on 3D confocal stacks. Number of cells were manually quantified using the Cell Counter Fiji plugin (*77*).

### Hybridization chain reaction and immunofluorescence

First, *emilin3a*:GFP, *emilin3a*:GFP-p2a-*her6*, *emilin3a*:GFP-p2a-*her9* or *emilin3a*:mScarlet-p2a-*jag1a* constructs were injected at the one cell stage and fish were fixed at 20-22 hpf. Hybridization chain reaction (Molecular Instruments) was performed following manufacturer instructions. *her6*, *her9*, *jag1a*, *jag1b* and *notch2* probes were produced by Molecular Instruments as 20 probe set sizes.

### Single cell RNA-Seq Analysis

Single cell RNA-Seq data was obtained from Wagner *et al*., 2018 (*35*). We filtered the raw data and selected the cells labelled as notochord in the original publication, and analyzed using the Scanpy v1.4.4 (*78*) python package. UMAP coordinates were calculated using normalized non-logarithmically transformed values and the scanpy.pp.neighbors function with n_neighbors = 20 and n_pcs = 5 parameter values. log(UMI+1) values were represented in the UMAP plots, where log represents natural logarithm. Boxplots and heatmaps were generated using the seaborn python package.

*emilin3a* was found as the gene with the best balance between notochord enrichment and high expression levels. We did this by selecting the gene with the highest score according to this equation:

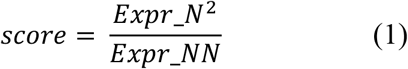

where *Expr*_*N* represents the average of normalized UMIs for each gene across all notochord cells at 18 hpf, and *Expr*_*NN* represents the analogous values for the non-notochord cells at the same stage. Genes with the highest score are shown in Table S3.

**Table S3.**
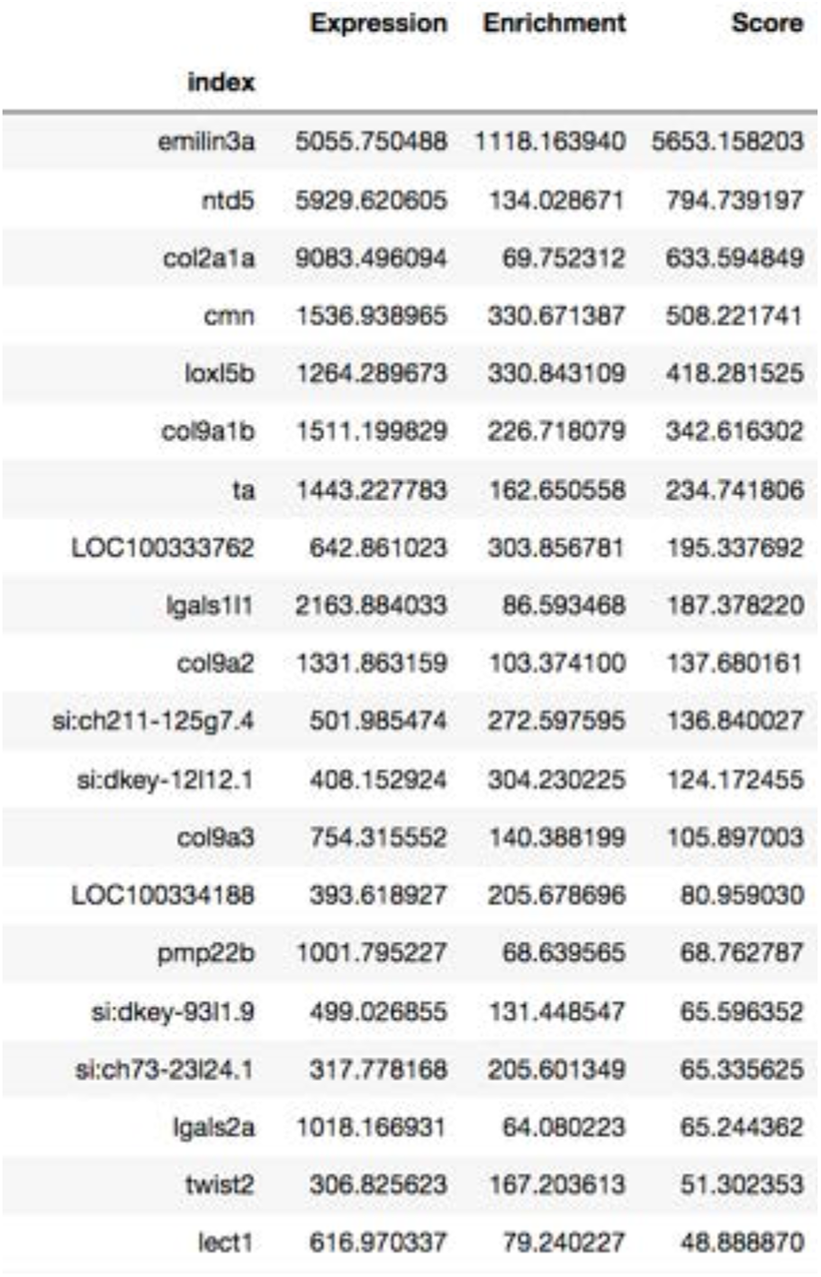
Genes with a highest score for specificity and expression levels in the notochord at 18 hpf. Expression: Average expression in Notochord cells (normalized UMIs per million). Enrichment: Average expression in notochord cells divided by average expression in the rest of the cells in the fish at 18 hpf. Score: Expression multiplied by enrichment (equivalent to the equation described above).

### Electron Microscopy

For EM imaging, samples were chemically fixed by immersing them in 2.5% glutaraldehyde and 4% paraformaldehyde in 0.1M PHEM buffer. Sections were post-stained with uranyl acetate for 5 minutes and with lead citrate for 2 minutes. The overall EM protocol is similar to previously reported (*79*).

### Microscopy

Zebrafish embryos were embedded in 0.6% agarose low gelling temperature (A0701, Sigma) with 0.16 mg ml−1 Tricaine in E3 medium. For imaging embryos between 18 and 24 hpf, agarose covering the tail was removed to allow freely development of their tail. Imaging was performed with a Zeiss LSM880 laser scanning confocal microscope, using a 40x/1.1NA water-immersion objective.

### Adaptive Feedback microscopy workflow

The adaptive feedback microscopy workflow was set up on Zeiss LSM880 AiryScan Fast microscope. Automated image analysis and definition of high-zoom tile positions was implemented as a Fiji plugin using previously developed AutoMicTools library (https://git.embl.de/halavaty/AutoMicTools). MyPic VBA macro (*80*) was used as a communication interface between the Fiji plugin and ZenBlack software controlling the microscope.

Both low-zoom and high-zoom images were acquired using AiryFast modality to enable time resolution of 5 minutes. 488nm line of the Argon laser was used for excitation, fluorescent signal was detected using 499-553 nm emission filter. Low-zoom images were acquired using lowest possible zoom and rectangular tilescan in the total area 991 by 673 μm with the pixel size 0.835 μm and spacing between slices 5 μm. Each high-zoom tile was acquired in the field of view 83.72 by 83.72 μm with the pixel size 0.108 μm and spacing between slices 2.5 μm. Collected high-zoom tiles were stitched in Fiji using BigStitcher plugin (*81*) and custom Jython scripts. To show the same region of the notochord independently on the move of the developing zebrafish, we used a custom-made Fiji Macro where the region of interest was manually selected every 10 frames, and interpolated for the rest of the timepoints.

To show the same region of the notochord independently on the move of the developing embryo, we used a custom-made Fiji Macro where the region of interest was manually selected every 10 frames, and the region of interest interpolated for the rest of the timepoints.

### Image analysis

Python 3.7.4 was used for image analysis. First, the intensities of each of the channels was normalized between 0 and 1. Then, a gaussian filter was applied to the channel. This was done using the filters.gaussian_filter function of scipy.ndimage package, with a sigma value equal to 3. Then, both adaptive and global single-value segmentation were applied to the GFP channel. For the global single-value segmentation, the value was chosen automatically for each image as 1.5 times the median intensity of the GFP channel. To generate the adaptive segmentation, we calculated the local mean using as a kernel a uniform circle of 120 pixel diameter, and the rank.mean function of the skimage.filters package. Only the pixels that with a higher value than both the global and the adaptive thresholds were considered for further analysis (Segmentation 1).

To define the GFP-positive cells, we filled holes in the cells by applying a 5-iteration binary dilation followed by a 9-interation binary erosion (scipy.ndimage python package). A higher erosion than dilation was applied to avoid defining as GFP-positive cells the pixels in the boundaries between cells. Only objects with an area of 3500 squared pixels were defined as cells and considered for further analysis (Segmentation 2).

The neighborhood of GFP cells was defined as follows. We first applied an 8-pixel binary dilation of 8 pixels to the GFP cells as defined in ‘Segmentation 1’ to define the boundary between cells. We then applied a 25-pixel binary dilation to define the neighboring cells. The region generated by the 25-pixel dilatation is the region that we considered as ‘neighboring cells’ (Segmentation 3).

To determine the relative intensity inside the ‘GFP-positive cells’ or the ‘neighboring to GFP cells’ we manually selected the notochord region, and we only considered the pixels inside the manually selected region. Then, the measured the mean value of the different mRNA signals inside the selected cells relative to the value of all the notochord.

In all the analyzed images, the stepsize is 63.7 nm/pixel. Plots were generated using boxplot and swarmplot functions of the seaborn python package.

### Statistical analysis

Statistical analysis was performed using the scipy.stats python package. The specific statistical test used, including sample size and the p-values are indicated in the figures and figure legends.

## DESCRIPTION OF THE THEORETICAL MODEL

### Lateral induction model

The lateral induction model was defined as a two-component system, Ligand (L) and Notch Intracellular Domain (NICD, represented as I in the equations). Notch-Ligand interaction in adjacent cells triggers the release of NICD following an increasing Hill function. NICD activates the expression of the ligand in its own cell following an increasing Hill function. The equations that describe the model are:

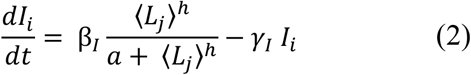

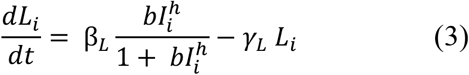

*L_i_* and *I_i_* are the average concentrations of Ligand and NICD inside the cells, respectively. 〈*L_j_*〉 is the average concentration of Ligand in each of the neighboring cells. β*_I_* and β*_L_* are the production rates of ligand and receptor, respectively. *γ_L_* and *γ_I_* are the degradation rates of Ligand and NICD, respectively, *a* and *b* the affinities, and ℎ is the Hill coefficient.

### Lateral inhibition model

This model is based on (*13*) and is similar to the lateral induction, with the only difference that the lateral inhibition model assumes that NICD activates the expression of a repressor that in turn inhibits the expression of the ligand. For this reason, the production of ligand is represented as an inhibitory Hill function.

The equations that describe the system are:

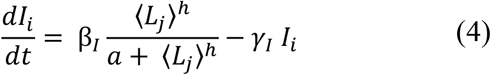

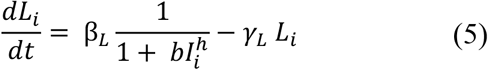

### Lateral inhibition model with mutual inhibition

The equations that describe this model are based on (*20, 21*)

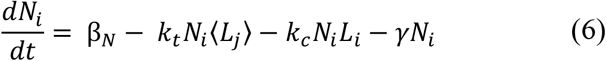

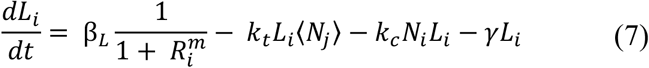

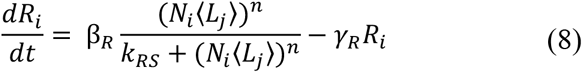

*N_i_*, *L_i_* and *R_i_* are the average concentrations of Notch Receptor, Ligand and Repressor inside the cells, respectively. 〈*L_j_*〉 and 〈*N_j_*〉 are the average concentrations of ligand in the neighboring cells. β*_N_,* β*_L_* β*_R_* are the production rates of Notch Receptor, Ligand and Repressor, respectively. γ and γ*_R_* are the degradation rates of Notch Receptor and Ligand/Repressor, respectively. K*_RS_* is the affinity, and *n*, *m* are the Hill coefficients for the different interactions. *k_t_* and *k_c_* are the interaction strength between ligand and receptor in *cis* and *trans*, respectively. These two constants are referred as *K_cis_*.and *K_trans_* in the manuscript.

### Simulations

All the visual simulations were generated by solving the equations using the Euler method with a step set to 0.01. Simulations were initialized with random values uniformly distributed between 0 and 0.1. To avoid boundary effects, we run simulations on a 100-cell array, where only the 20 central cells are displayed, while the 40 cells in each side buffer the boundary effect.

### Linear stability analysis

Linear stability analysis was done as previously described (*20*). A prerequisite for pattern formation is the instability of the homogenous steady state (*N*\*, *N*\*, *L*\*), where every cell has the same value of *N_i_*, *N_i_* and *L_i_*. We first calculated the homogeneous steady state by making *N_i_* and *N_j_* equal to *N*\*, *N_i_* and *N_j_* equal to *N*\*, and *L_i_* equal to *R*\*, and solving the following system of equations (*20*):

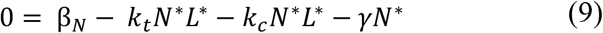

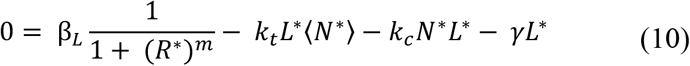

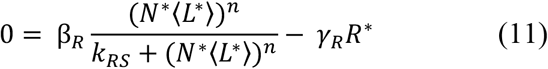

We solved these equations for the *L*\*, *N*\* and *N*\* using the fsolve function of the scipy.optimize python package.

The stability analysis requires the computation of the Jacobian matrix, that according to Othmer and Scriven (*82*) can be expressed as *J* = *I_k_* ⊗ H + M ⊗ B, where *I_k_* is the *k x k* identity matrix, *k* is the number of cells, ⊗ represents the tensor product, 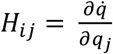 is the change in production of species *i* for a change in species *j* in the same cell, 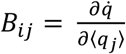 is the change in production of species *i* for a change in species *j* in a neighboring cell, and *M* is the connectivity matrix defined as

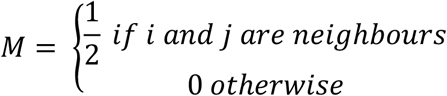

In the specific case of our model, where cells are arranged unidimensionally, *i* and *j* are neighbors when |*i* − *j*| = 1.

The eigenvalues of *J* Jacobian matrix are the eigenvalues of the various matrices *H* + *q_k_B*H, where *q_k_* are the eigenvalues of the connectivity matrix *M*. For our particular *M* matrix, *q_k_* values are always higher or equal to – 1, meaning that we only need to compute an eigenvalue for the extreme case *q_k_* = −1 to determine the highest eigenvalue (known as the Maximum Lyapunov Exponent, MLE) has a positive real part.

Following this strategy, we computed the MLE value for a grid of *k_c_* and *k_t_* values logarithmically spaced between 0.001 and 100.

### Parameter values

|  |  |
| --- | --- |
| Figs 1 and S2 | $a = 0.1$<br>$b = 10$<br>$h = 2$<br>$\gamma_I = 1$<br>$\gamma_L = 1$<br>$\beta_I = 1$<br>$\beta_L = 1$ |
| Figs 5 and S8 | $m = 1$<br>$n = 1$<br>$k_c = 0.001 \text{ to } 100$<br>$k_t = 0.001 \text{ to } 100$<br>$\gamma = 1$<br>$\gamma_R = 1$<br>$k_{RS} = 1$<br>$\beta_N = 2$<br>$\beta_L = 10$<br>$\beta_R = 10$ |

## AUTHOR CONTRIBUTIONS

Conceptualization: H.S-I and A.D-M. Methodology: H.S-I and A. H. Investigation: H.S-I. Visualization: H.S-I and A. H. Software: H.S-I and A. H. Supervision: A.D-M. Writing – original draft: H.S-I and A.D-M. Writing – Review & Editing: all authors Funding Acquisition: H.S-I and A.D-M.

## ACKNOWLEDGEMENTS

We thank Anna Erzberger, Aissam Ikmi and Stefano de Renzis for critical reading of the manuscript. We thank Jonas Hartmann for discussion on the project and training on image analysis. We are grateful to the EMBL EM core facility (EMCF), and in particular to Rachel Mellwig and Yannick Schwab, for the EM experiments. We thank the EMBL Fish Facility, and in particular to Sabine Görgens. We thank the EMBL Advanced Light Microscopy Facility, and especially Christian Tischer and Stefan Terjung for image analysis and microscopy support. We thank Alexander Ernst for the development of the custom-made ImageJ macro used to generate some of the movies of the paper. We thank ZFIN for making zebrafish information easily accessible (*69*). We thank the Life Science Editors for editorial support.

## SUPPLEMENTARY INFORMATION

**Figure S1.**
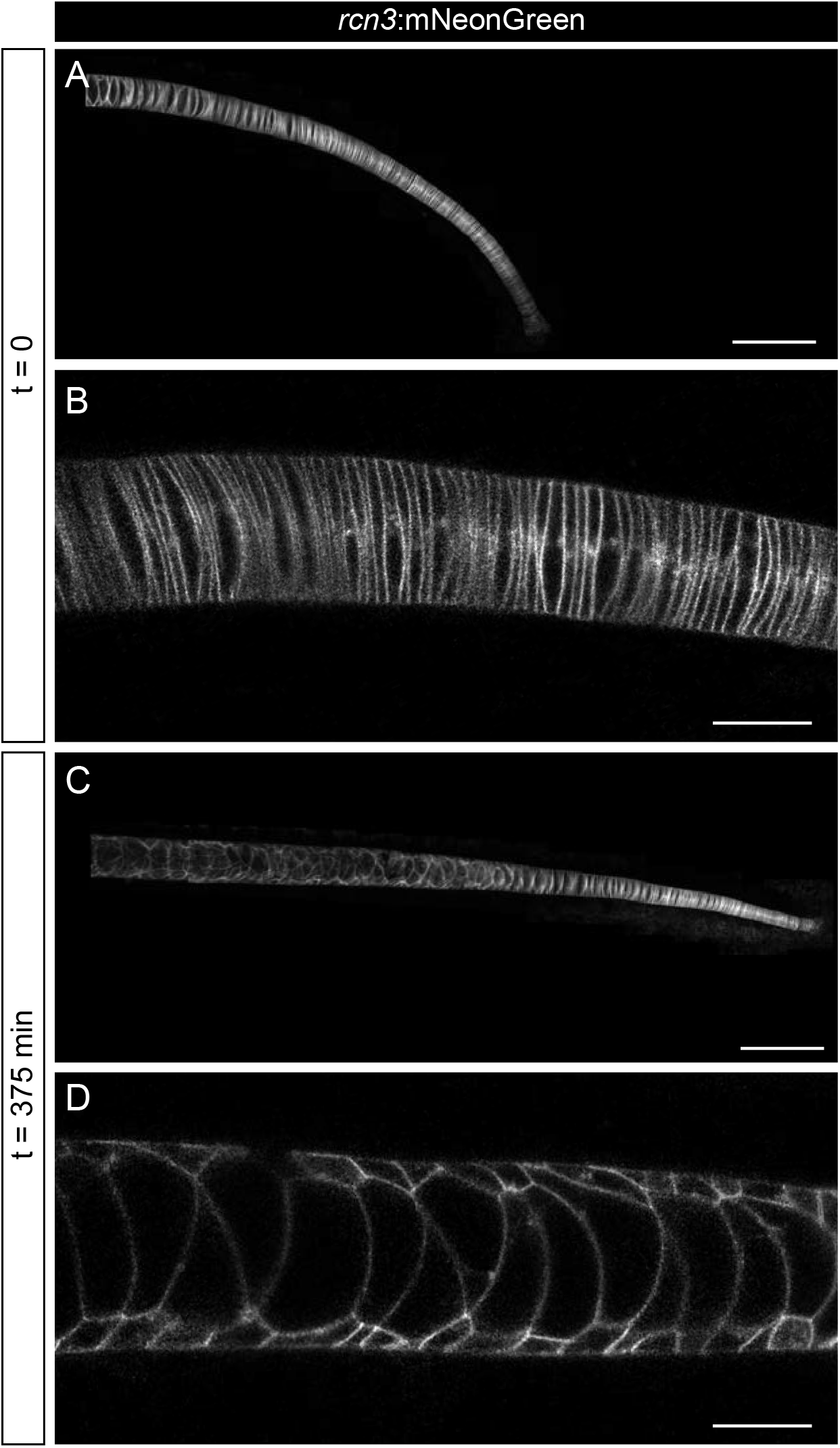
*In vivo* imaging of notochord development. (**A**-**D**) *In vivo* time-lapse imaging of zebrafish notochords starting acquisition at 22 hpf using the *rcn3*:mNeonGreen transgenic line. Acquisition was based on a feedback microscopy protocol, where low quality images were first acquired and then analyzed at the time of acquisition to perform high zoom tile scan imaging only in the notochord cells. (A, C) show maximum projection of Airyscan confocal notochord reconstructions. (B, D) show zoomed images of single Airyscan confocal optical sections magnified from (A, C), respectively. Scale bars, 100 μm (A, C), and 20 μm (B, D).

**Figure S2.**
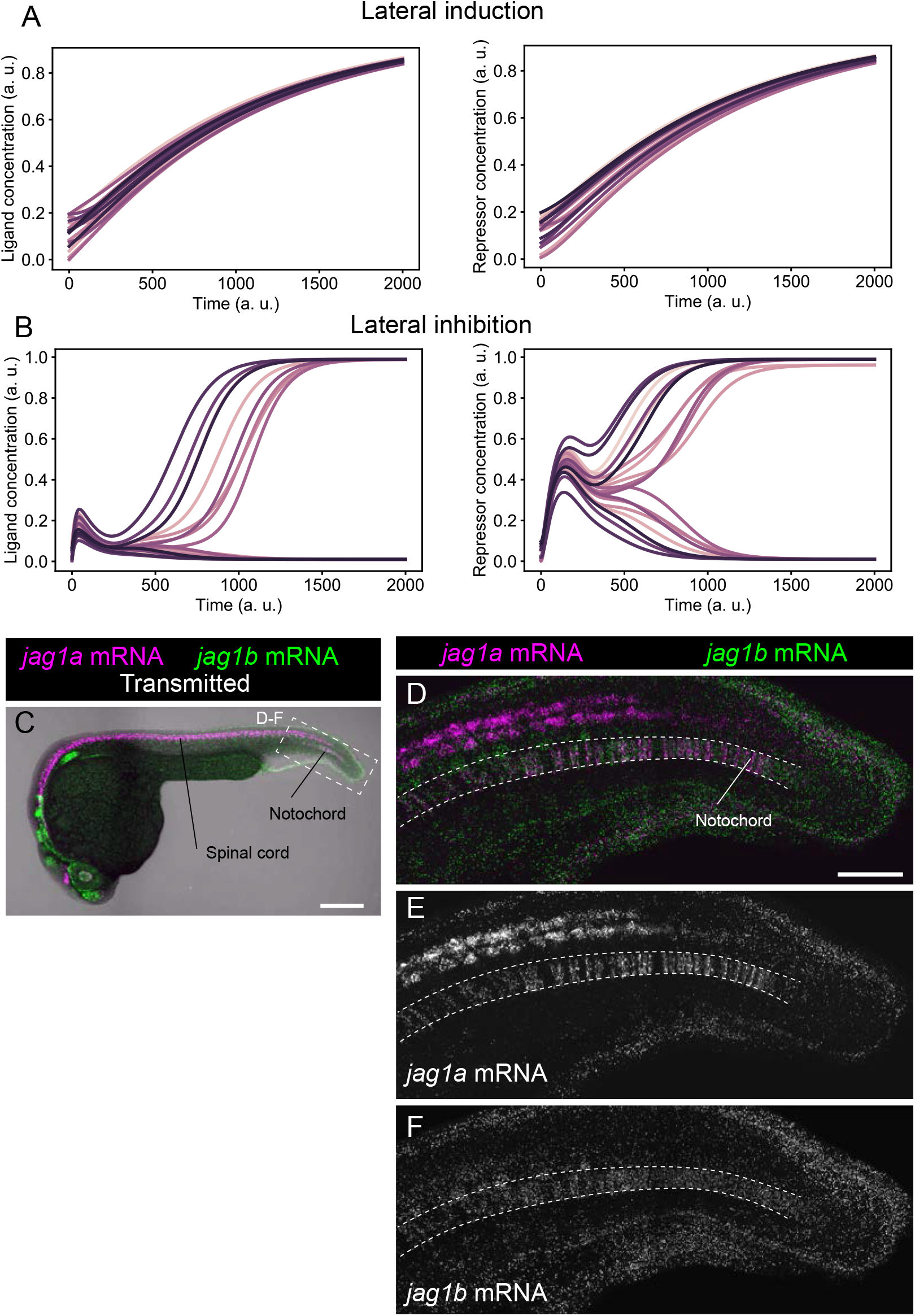
Lateral induction and lateral inhibition simulations, and mRNA expression pattern of *jag1a* and *jag1b*. (**A** and **B**) Representative simulations of NICD and Ligand molecules in each cell for the lateral induction (A) and lateral inhibition (B) models. Each line represents a different cell. Related to models and simulations shown in Figures 1E and 1F. (**C**) Confocal projections of 24-hpf zebrafish stained with *in situ* HCR probes against *jag1a* (magenta) and *jag1b* (green). (**D**-**F**) Airyscan confocal projections at a higher magnification of boxed region in (C). n = 5 fish. Scale bars, 200 μm (C), and 50 μm (D).

**Figure S3.**
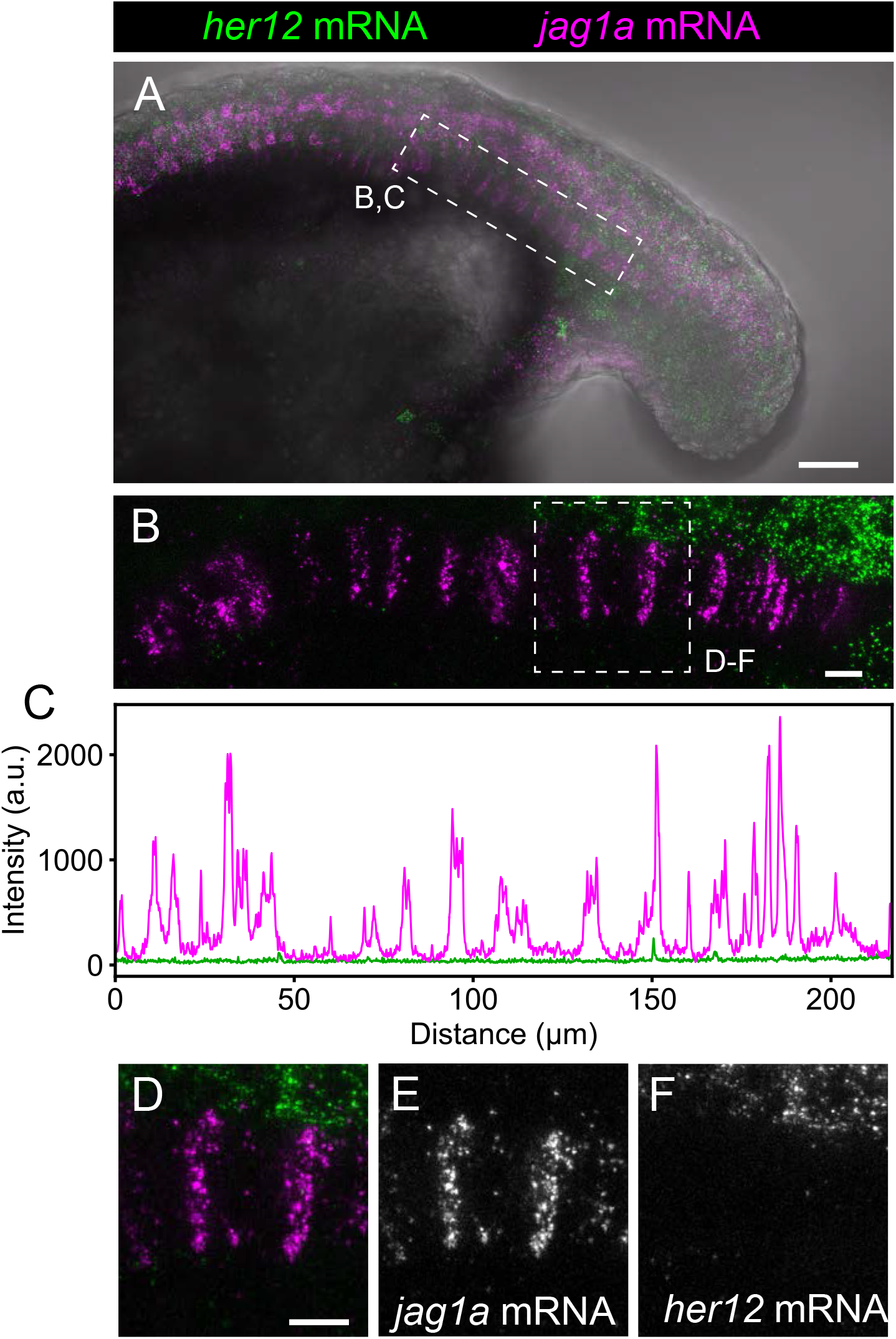
*her12* expression is not detected in the notochord. (**A**) Projection of confocal optical sections of 18 hpf zebrafish stained with in situ HCR probes against *her12* (green) and *jag1a* (magenta). Transmitted light is shown in gray scale. (**B**) Maximal projection of confocal Airyscan optical sections of the boxed area in (B)**. C**, Intensity profile of *her6* (green) and *jag1a* (magenta) along the notochord based on in situ HCR shown in (B). (**D**-**F**) Magnified views of boxed area in (B) n = 8. Scale bars, 50 μm (A) 20 μm (B, D).

**Figure S4.**
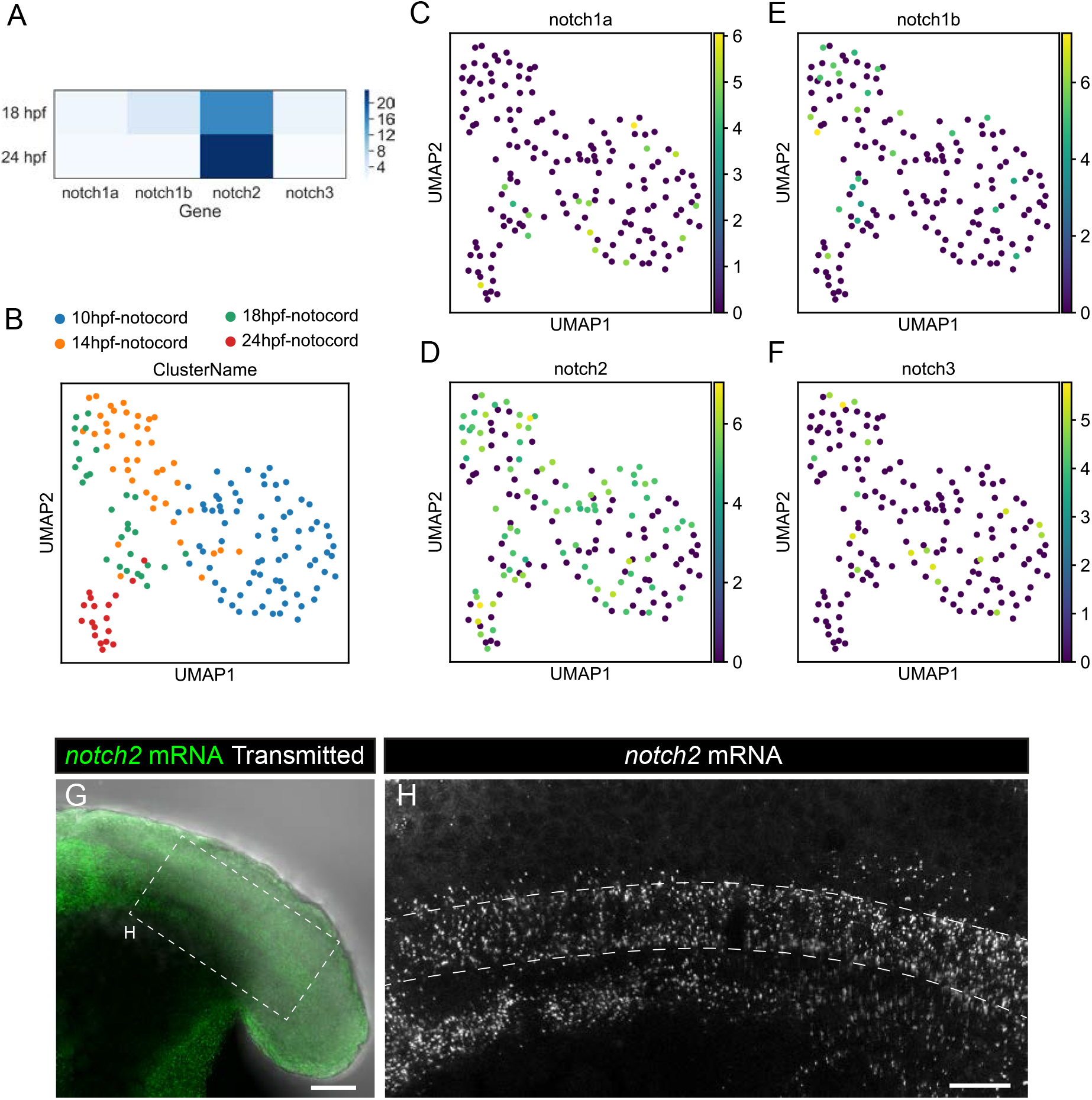
Expression of Notch receptors. (**A**) Heatmap showing the expression levels of the Notch receptor genes. Values represent average normalized UMIs in all notochord cells at 18 and 24 hpf. (**B**-**F**) UMAP plots showing notochord cells at 10, 14, 18 and 24 hpf. Cells are labeled depending on the developmental stage (A) or using a logarithmic color scale of of *jag1a* (B), *her6* (C) and *her9* (D) normalized expression. (**G**) Projection of confocal optical sections of 18 hpf zebrafish stained with in situ HCR probe against *notch2* (green). Transmitted light is shown in gray scale, n = 11. (**H**) Projection of confocal Airyscan optical sections of the boxed area in (G). Scale bars, 50 (G) 20 μm (H).

**Figure S5.**
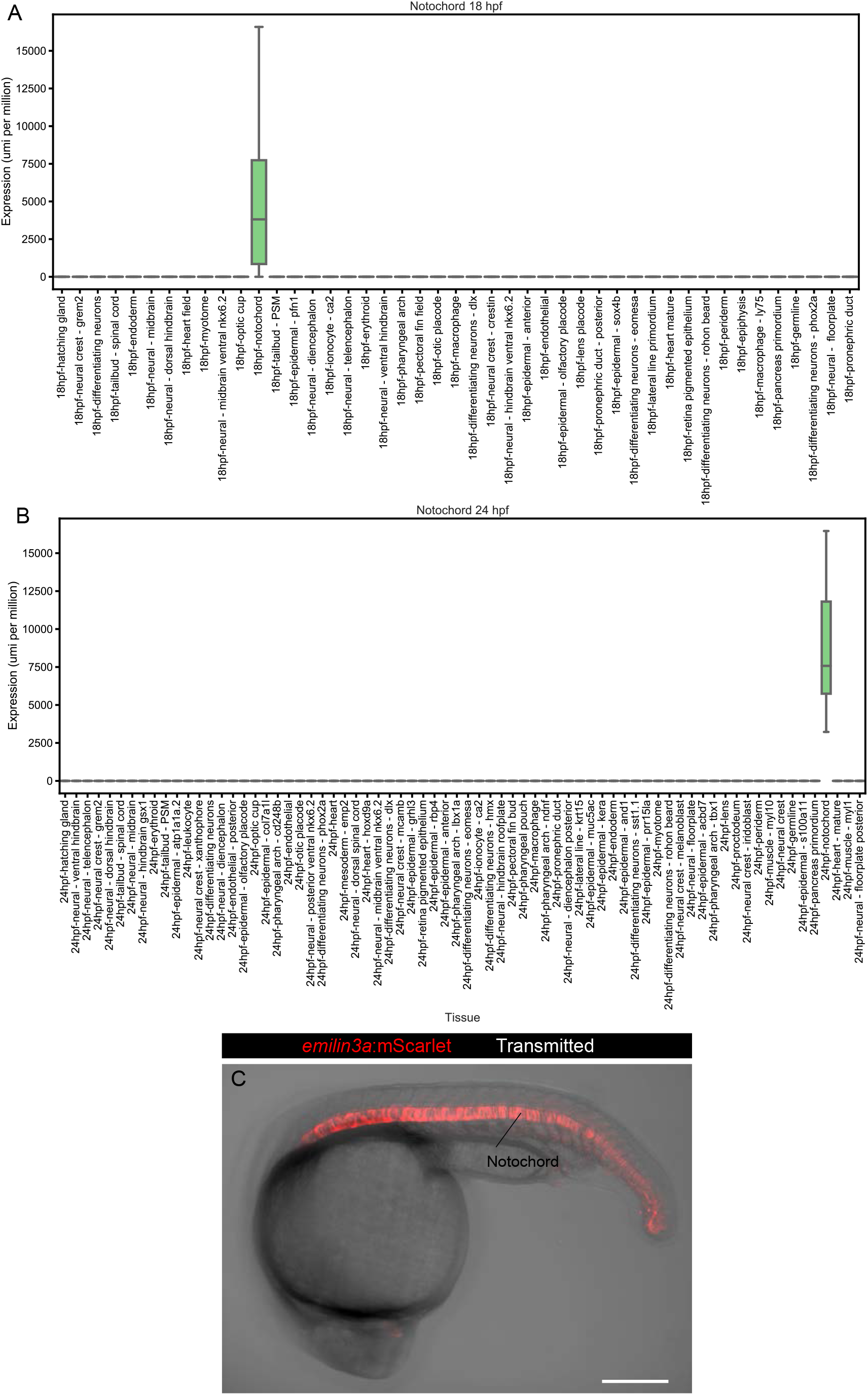
*emilin3a*-5kb promoter drives expression to the notochord. (**A** and **B**), Boxplot representing *emilin3a* expression in different tissues at 18 (A) and 24 hpf (B) according to single-cell transcriptomics. (C) Maximum projection of optical confocal sections of an *emilin3a*:mScarlet transgenic line at 22 hpf. Transmitted light is shown in gray scale. Scale bar, 200 μm.

**Figure S6.**
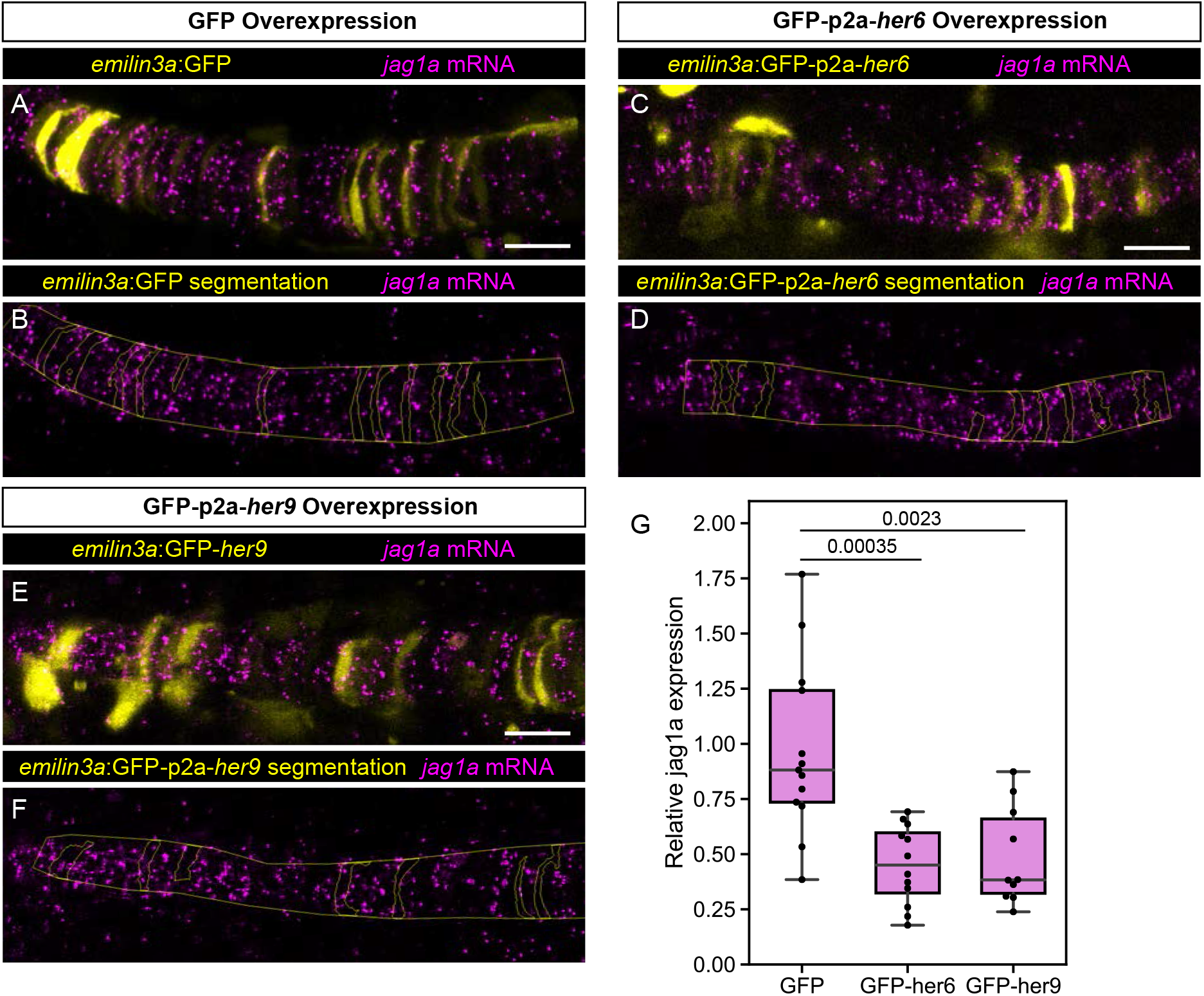
*jag1a* mRNA expression upon *her6* or *her9* overexpression. (**A** - **F**) Airyscan confocal optical sections of fixed 22 hpf transgenic injected with *emilin3a*:GFP (A and B), *emilin3a*:GFP-p2a-*her6* (C and D) or *emilin3a*:GFP-p2a-*her9* (E and F) constructs. GFP was detected by antibody staining and *her6* and *her9* mRNA by *in situ* HCR in whole mount embryos. (B, D, F) show the boundary of GFP segmentation in A, C and E, respectively, and manual outline of the notochord. (**G**) Quantification of *jag1a* mRNA intensity inside GFP-positive cells segmented as exemplified in A-F. Each point represents an individual fish.

**Figure S7.**
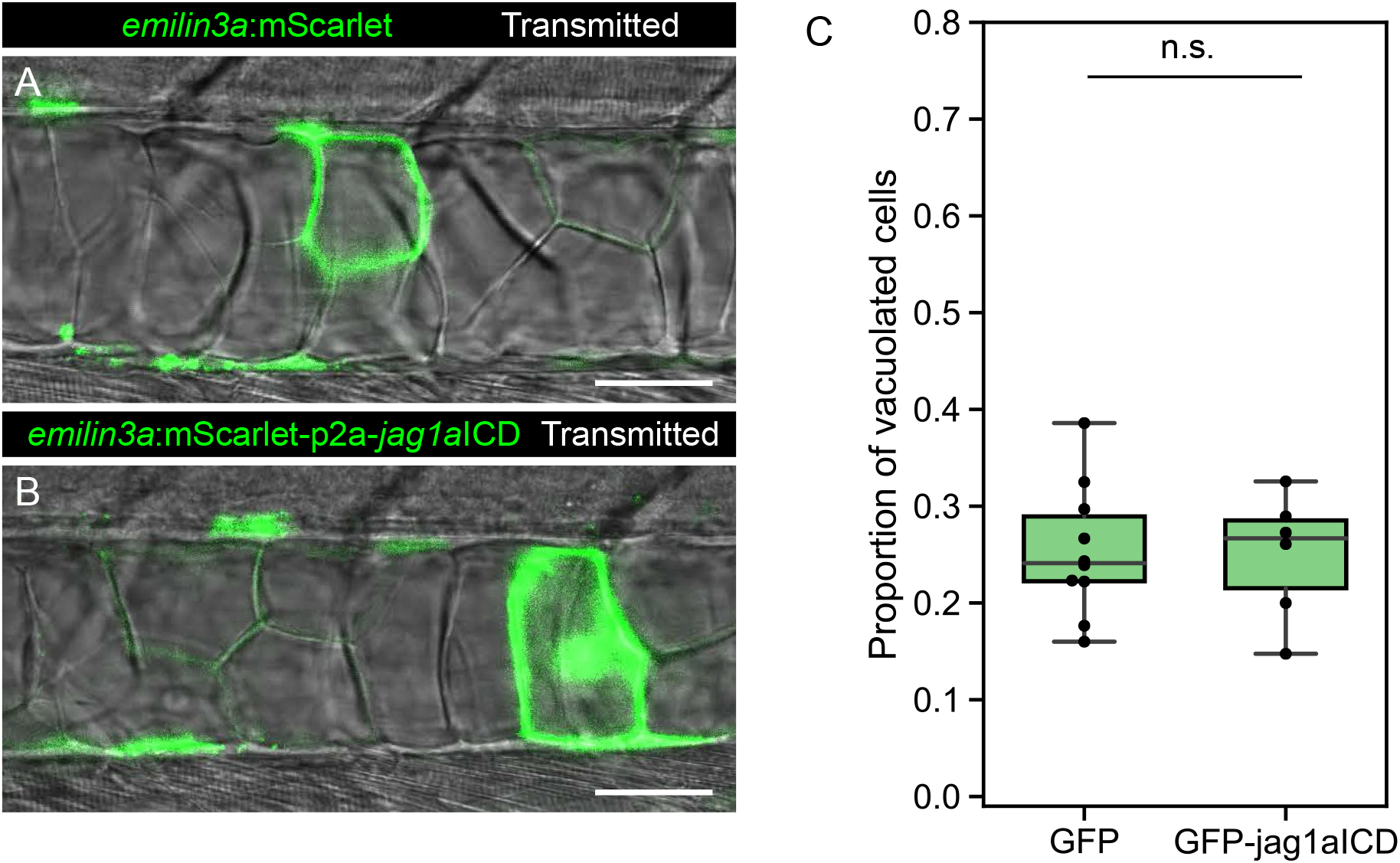
Jag1a intracellular domain does not have an effect on notochord cell fate. (**A**-**B**), Optical sections of 2 dpf zebrafish that were injected with the *emilin3a*:GFP or *emilin3a*:GFP-p2a-*JICD* constructs. (**C**) Proportion of vacuolated cells at 2 dpf are shown. Each point represents an independent fish quantified from on z-stack confocal acquisitions. (n.s.) p-value > 0.05 by two-tailed t-test. Scale bars, 50 μm.

**Figure S8.**
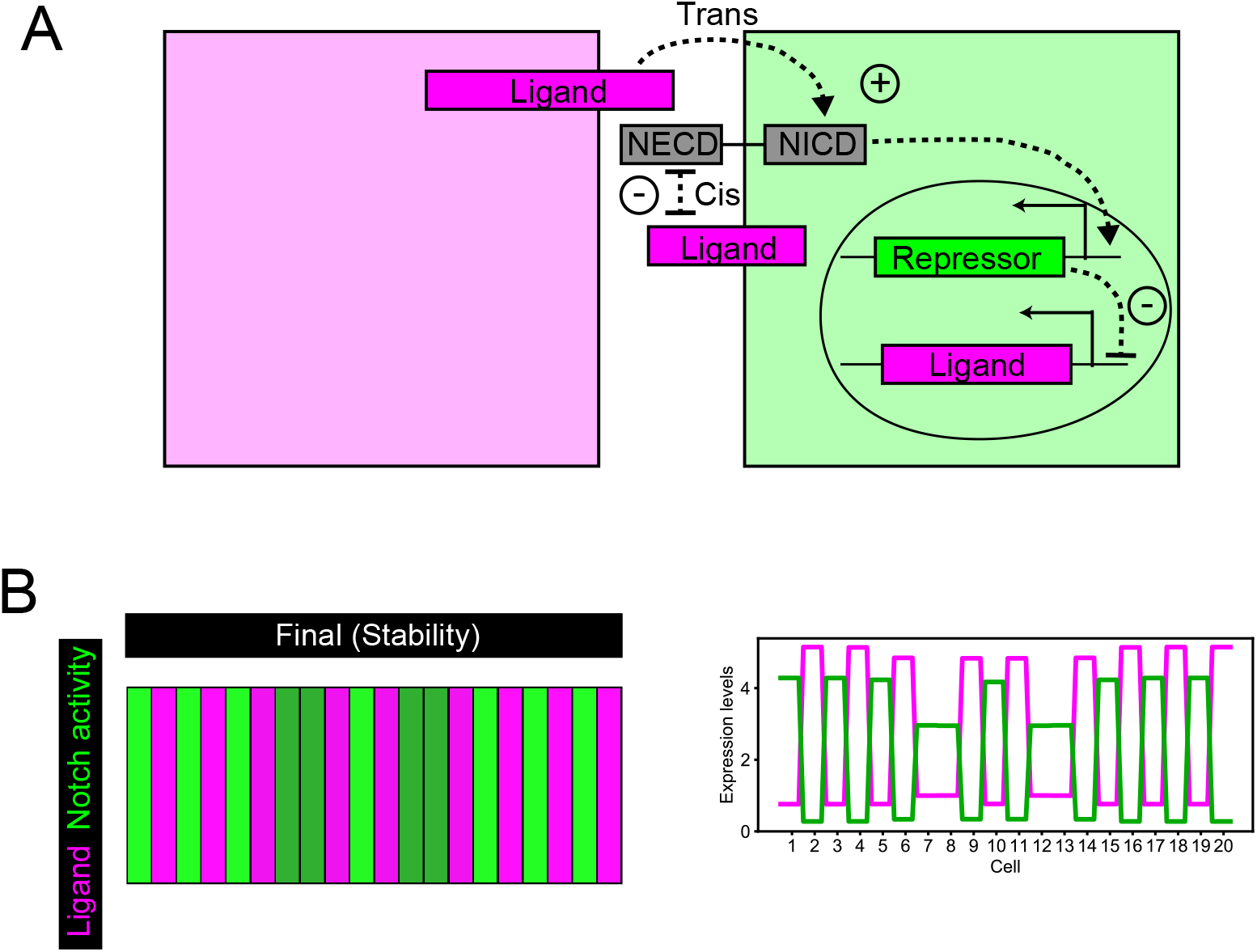
(**A**) **Model of lateral inhibition including cis interactions.** Schematic representation of a lateral inhibition model with mutual inhibition. Ligand binding to Notch receptor releases NICD, that activates Repressor expression. The repressor inhibits Ligand expression. In addition, when Notch receptor and Ligand are present in the same cell, they mutually inhibit to each other. (**B**) Representative simulation of the model shown in (A).

## SUPPLEMENTARY FILES

**Movie S1.** Maximal projection of the zebrafish notochord optical planes acquired using the feedback microscopy protocol to optimize quality of the region of interest.

**Movie S2.** Selected plane from Movie S1 stabilizing and magnifying a specific region of the notochord.

**Movie S3.** Time lapse optical section of notochord cells using the tp1:GFP; *jag1a*:mScarlet double transgenic line.

**Data S1**. mScarlet FRT kan FRT plasmid

**Data S2**. emilin3a mScarlet

**Data S3**. rcn3 lyn mNeonGreen

**Data S4**. emilin3a GFP

**Data S5**. emilin3a GFP-p2a-her6

**Data S6**. emilin3a GFP-p2a-her9

**Data S7**. emilin3a mScarlet-p2a-JICD

